# Optimizing Oxylipin Analysis with Liquid Chromatography Mass Spectrometry through Bio-Inert Systems and Ion Funnel Adjustments

**DOI:** 10.64898/2026.07.27.741082

**Authors:** John T. Shuster, Lingshuang Wu, Jericha Mill, Michaela Morhaus, Nina Yu Fan, Fernando Tobias, Dominique Baldwin, Matthew D. Bruss, Liam D. Hurley, Michelle E. Kimple, Adam R. Konopka, Judith A. Simcox

## Abstract

Oxylipins are potent signaling lipids that affect inflammation, vascular tone, and metabolism, making them relevant in many diseases. Oxylipins are measured with liquid chromatography-mass spectrometry (LC-MS), but challenges in quantification arise due to low abundance and rapid degradation. In this study, we optimize LC-MS methods to improve the quantification of oxylipins in human plasma, given growing interest in oxylipins and their impact on clinical research. Plasma samples were obtained from healthy participants, and oxylipins were concentrated by solid-phase extraction. Quantification was performed with a reversed-phase targeted LC-MS/MS method on an Agilent 6495D triple quadrupole with transitions for 248 oxylipin species. Given the rapid degradation of oxylipins with bio-reactive surfaces, we compared both standard and bio-inert (Altura) columns, standard and bio-inert LC setups, and ion funnel voltage. By decreasing voltages applied to ions inside the ion funnel, signal was increased for most oxylipin species while peak quality was maintained. Fully bio-inert setups quantified more compounds and had increased levels of some compounds, but fewer epoxyoctadecadienoic acid (EpODE) species. To explore this further, analytical-grade alpha-linolenic acid (ALA), the direct precursor of EpODEs, was injected and observed to form EpODEs within the instrumentation when using stainless steel columns. These data suggest that oxylipins benefit from fully bio-inert systems and optimized pre-mass analyzer parameters. The stainless-steel components of the column may also contribute to epoxidation reactions of polyunsaturated fatty acids (PUFAs), generating oxylipin species during analysis. Finally, various human sample matrices were assessed for compatibility with oxylipin quantification, including granulocytes, mononuclear cells, erythrocytes, skeletal muscle, and THP-1 cells, a human-derived monocyte cell line.

## 1. Introduction

Oxylipins are a class of signaling lipids with hundreds of species that regulate numerous facets of disease, including inflammation, mitochondrial abundance, vascular remodeling, and energy expenditure.^1–3^ Oxylipins are primarily produced from polyunsaturated fatty acids (PUFAs) by a variety of enzymes, including cyclooxygenase, lipoxygenase, and cytochrome P450 (CYP450) enzymes, as well as reactive oxygen species.^4^ **Figure 1** demonstrates the various PUFA precursors of oxylipins and which oxylipins they produce. Epoxide oxylipins are of particular importance and have been demonstrated to alter monocyte activation and differentiation in human-derived samples through activation of p38 MAPK.^5^ Epoxides are also associated with the development of type 1 diabetes in at-risk children, as well as protection against hypoxia and reperfusion injury.^6,7^

**Figure 1:**
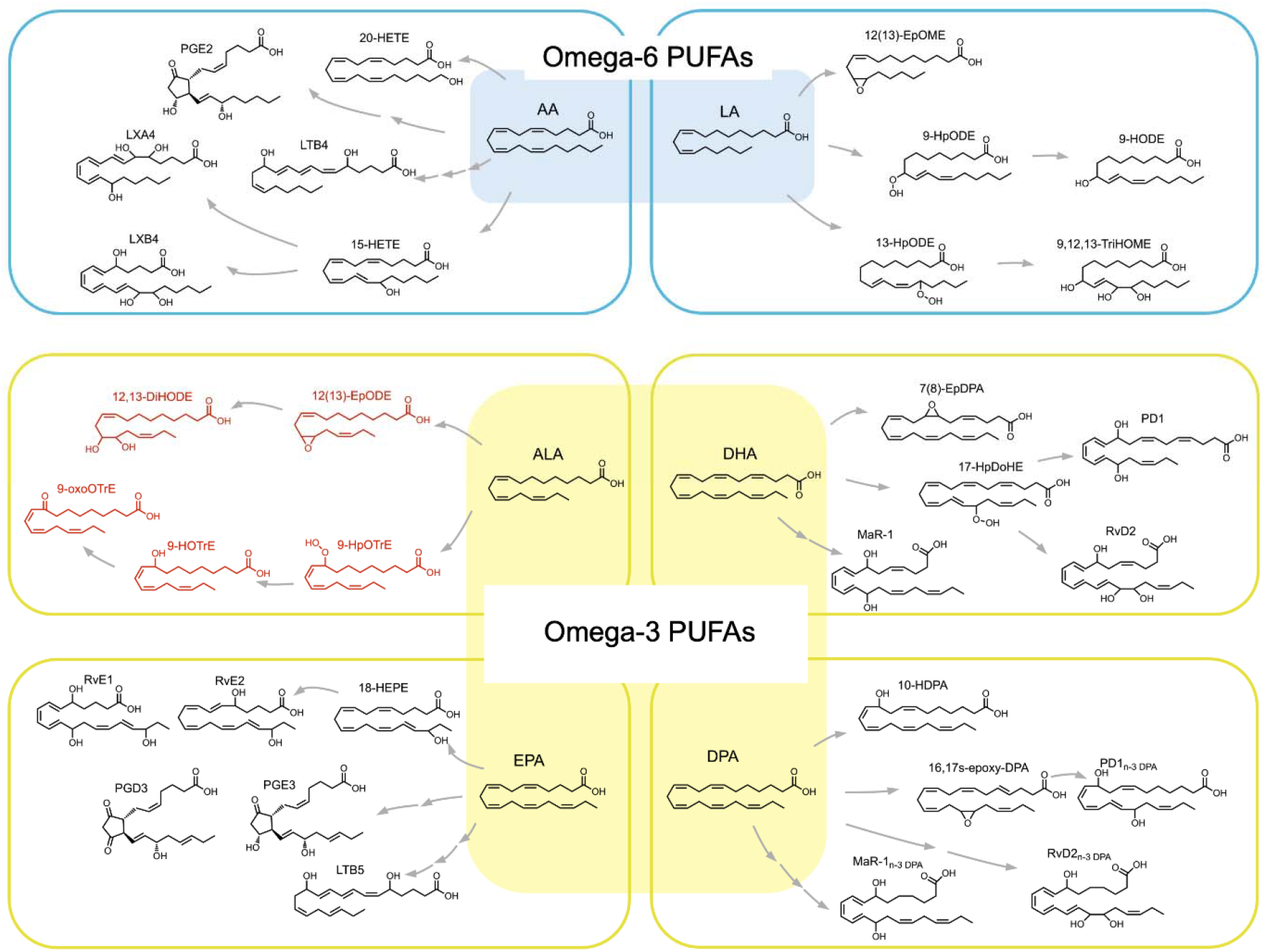
Omega-6 and Omega-3 PUFAs Generate Distinct Oxylipin Families. Chart showing oxylipin families grouped by precursor PUFA. Blue denotes omega-6 PUFAs and omega-6–derived oxylipins. Yellow denotes omega-3 PUFAs and omega-3 derived oxylipins. Red indicates ALA-derived products that we show can be formed by oxidation in non-bio-inert LC systems and columns.

Oxylipins, due to their status as signaling molecules, are present at low abundance and have a short half-life.^8,9^ In this work, “oxylipin” refers to oxygenated metabolites of monounsaturated fatty acids (MUFAs) and PUFAs, regardless of chain length or origin.^9^ To overcome the low abundance and stability, solid-phase extraction is used to purify and concentrate oxylipins from tissue or plasma into a prepared sample. Careful method design is also vital to minimize ion loss or in-source fragmentation, while ensuring all oxylipins reported have actually been detected. The field of oxylipin analysis is at a turning point, with some reports of species detected lacking adequate chromatography or criteria to claim detection or quantification.^10^

One challenge to overcome with LC-MS is the metal adsorption interactions of oxylipin carboxylic acid groups with the stainless steel components of the chromatography system.^11^ In traditional high-performance liquid chromatography (HPLC) systems, passivation can be used to decrease stainless steel interactions with analytes.^12^ This is typically done by pretreatment of the entire LC flow path with phosphoric or citric acid, which has been shown to decrease peak tailing and increase peak height.^11,12^ Another technology frequently used in lipid LC-MS is the use of bio-inert systems, with previous studies reporting improvements in analysis of various lipid classes using fully bio-inert systems.^13^ These systems utilize a bio-inert HPLC system and chromatography columns, achieved through replacement of stainless steel with titanium, MP35N, or polyetheretherketone/proprietary coatings on stainless steel surfaces. This fully bio-inert system has been shown to improve certain lipid relative intensities and separation efficiency, but it has not yet been shown to have notable effects on oxylipins or increase detectable compounds.^13^ The field of oxylipin analysis does not yet utilize fully bio-inert chromatography systems as standard for the field.^9,13,14^ Bio-inert LC systems are occasionally used, but it is less common to observe bio-inert LC systems combined with bio-inert chromatography columns.

Another point of optimization is ion guiding parameters. After electrospray ionization, nebulized ions first enter the mass spectrometer, then pass through ion guides and into the mass analyzers, driven by electric forces and pressure differentials. Radiofrequency (RF) voltages are then applied to confine and direct the maximum number of ions into the mass analyzers.^15^ It has been previously demonstrated that there is a balance between using higher voltages during ion guiding to maximize the number of ions entering the mass analyzer and limiting pre-mass analyzer fragmentation through lowering RF voltages.^16^ For example, high RF amplitudes can fragment certain small perfluoroalkyl carboxylates (per- and polyfluoroalkyl substances, PFAS) by loss of CO2, with fragmentation approaching 100% for some compounds.^16^ Previous work on macromolecular assemblies has also shown that RF amplitude tuning in pre-mass analyzer ion optics can select for single precursor ion transmission, further reinforcing the importance of ion optics variables.^17^ Currently, there is no central guidance on the optimal ion guide RF voltage for oxylipins, as each compound may require individual RF voltages.^16^ This was of particular interest to us, as these parameters can be adjusted on Agilent systems. Overall, there are many factors in oxylipin LC-MS that present opportunities for optimization to better detect these low-abundance compounds.

Here, we show several improvements in oxylipin quantification using lower ion guide voltages and fully bio-inert chromatography systems. Lower ion guide voltages improved the relative intensity of low abundance oxylipin species, and bio-inert chromatography systems allowed for detection of more oxylipin species. We also show stainless steel chromatography columns may modify oxylipins and their precursors inside of the LC system. Finally, we demonstrate this method on a variety of human tissues including erythrocytes, peripheral blood mononuclear cells, granulocytes, skeletal muscle, and THP-1 cells.

## 2. Experiment

### Sample Generation

All studies were approved by the University of Wisconsin-Madison Institutional Review Board (IRB# 2023-0230). Blood samples were obtained from a healthy participant and collected in EDTA blood collection tubes on ice, as EDTA is preferred in oxylipin analysis due to interactions between lipids and other anticoagulants.^14,18^ Plasma was separated by centrifugation at 1,500 *× g* for 10 minutes at 4°C. Additional plasma was obtained from Innovative Research (IPLAK3E100ML-48663) for verification of results seen with the healthy plasma, which was generated and frozen at -80°C for 24 months before analysis. Samples were aliquoted and processed to limit freeze thaw cycles; healthy participant plasma was stored at -80°C for a maximum of 2 weeks. Skeletal muscle biopsy samples were obtained from the vastus lateralis under local anesthesia (1% xylocaine), dissected free of visible adipose and connective tissue, snap-frozen in liquid nitrogen, stored at -80°C, and homogenized prior to analysis. Human islets were obtained from the University of Wisconsin - Madison Human Islet Core. Islets were recovered in a 50:50 mixture of PIM(T) and PIM(R) media for at least 72 hours after isolation. Islets were handpicked into pellets of 100, washed with PBS, and spun down before excess PBS was removed and pellets were snap frozen in liquid nitrogen prior to extraction.

### Blood Fractionation

Blood fractions were obtained according to manufacturer protocol (Cytiva 17144002). Briefly, 3 mL of Ficoll-Paque Plus was added to a 15 mL tube, and 4 mL of 1:1 whole blood to PBS was overlaid; standards were added into select blood samples at this point. Samples were centrifuged at 400 *× g* for 30 minutes with the brake off. Fractionation blank and spiked-buffer blank samples of PBS with and without standards in place of blood were prepared as well. The plasma layer was drawn off, then the mononuclear, granulocyte, and erythrocyte layers were sequentially collected. The mononuclear cell layer was diluted with 5 volumes of sterile PBS and centrifuged at 400 *× g* for 30 minutes to wash the cells. The supernatant was removed, and cells were frozen until analysis. The granulocytes were suspended in 5 volumes of sterile PBS and centrifuged at 400 *× g* for 30 minutes to wash the cells. The supernatant was removed, and 10 volumes of red blood cell lysis buffer were added, and the sample was incubated at room temperature for 10 minutes. The cells were spun again at 400 *× g* for 30 minutes, and the supernatant was removed. Granulocytes, erythrocytes, and PBMCs were then frozen until analysis, for no more than 2 weeks. Antioxidant was not added for any steps in blood fractionation. All centrifugation steps for blood fractionation were done at 20 °C.

### Reagents

All internal standards were obtained from Cayman Chemical. Standards were added in equal quantities to a master mix (**Table 1**).

**Table 1:**
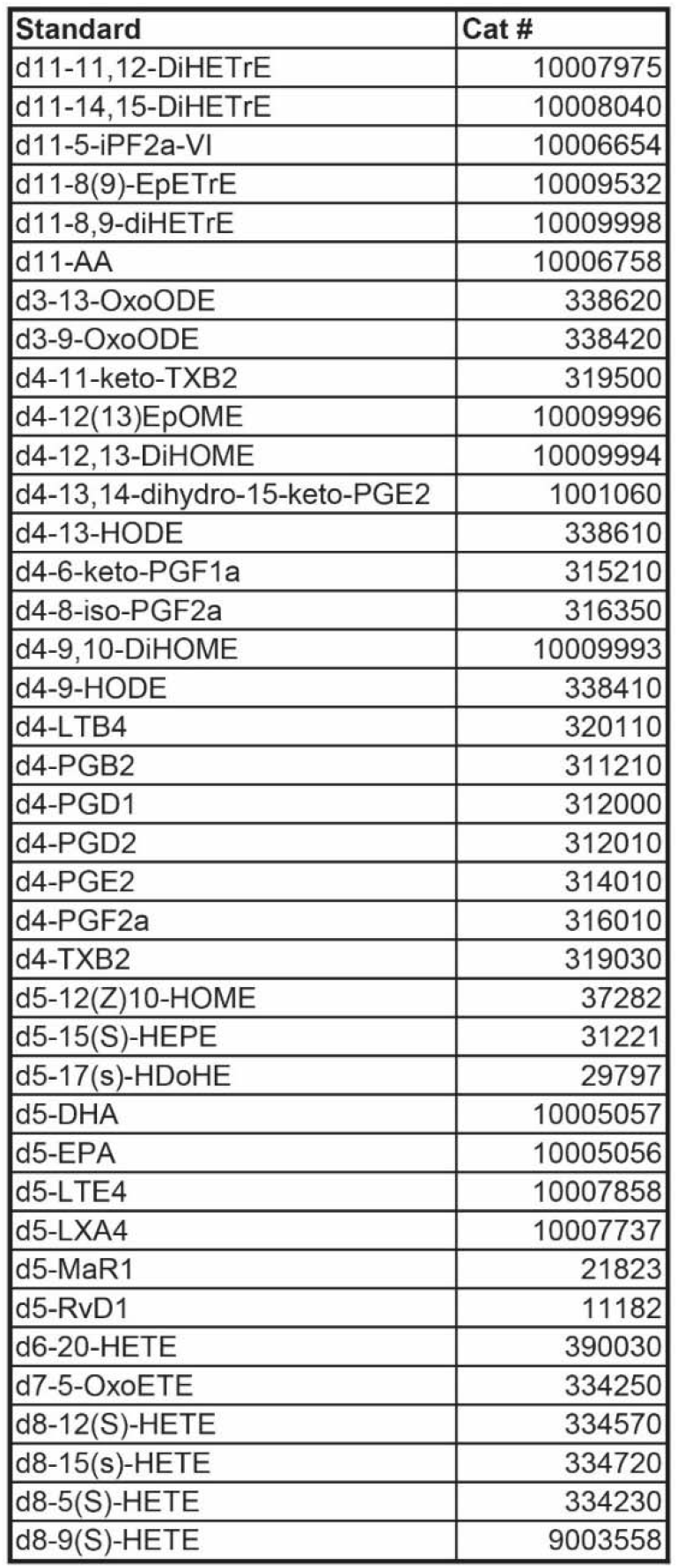
Internal Standards Added to Oxylipin Extractions. All internal standards were added in equal amounts to achieve a final concentration of 6 ng/mL in the oxylipin extract. All compounds were obtained from Cayman Chemical.

| Standard | Cat # |
| --- | --- |
| d11-11,12-DiHETrE | 10007975 |
| d11-14,15-DiHETrE | 10008040 |
| d11-5-iPF2a-VI | 10006654 |
| d11-8(9)-EpETrE | 10009532 |
| d11-8,9-diHETrE | 10009998 |
| d11-AA | 10006758 |
| d3-13-OxoODE | 338620 |
| d3-9-OxoODE | 338420 |
| d4-11-keto-TXB2 | 319500 |
| d4-12(13)EpOME | 10009996 |
| d4-12,13-DiHOME | 10009994 |
| d4-13,14-dihydro-15-keto-PGE2 | 1001060 |
| d4-13-HODE | 338610 |
| d4-6-keto-PGF1a | 315210 |
| d4-8-iso-PGF2a | 316350 |
| d4-9,10-DiHOME | 10009993 |
| d4-9-HODE | 338410 |
| d4-LTB4 | 320110 |
| d4-PGB2 | 311210 |
| d4-PGD1 | 312000 |
| d4-PGD2 | 312010 |
| d4-PGE2 | 314010 |
| d4-PGF2a | 316010 |
| d4-TXB2 | 319030 |
| d5-12(Z)10-HOME | 37282 |
| d5-15(S)-HEPE | 31221 |
| d5-17(s)-HDoHE | 29797 |
| d5-DHA | 10005057 |
| d5-EPA | 10005056 |
| d5-LTE4 | 10007858 |
| d5-LXA4 | 10007737 |
| d5-MaR1 | 21823 |
| d5-RvD1 | 11182 |
| d6-20-HETE | 390030 |
| d7-5-OxoETE | 334250 |
| d8-12(S)-HETE | 334570 |
| d8-15(s)-HETE | 334720 |
| d8-5(S)-HETE | 334230 |
| d8-9(S)-HETE | 9003558 |

### Oxylipin extraction

Lipid extraction from plasma was performed using solid-phase extraction. Oxylipin quantitation was achieved by using a master mix composed of 20/80 methanol/water (v/v) with 0.02 mg/ml butylated hydroxytoluene (BHT) and 6 ng/mL each of 39 deuterated oxylipin standards (**Table 1**). 250 μL plasma samples were transferred into 1.5 mL tubes and vortexed with 750 μL master mix. Samples were incubated on ice for 5 minutes and intermittently vortexed, then centrifuged at 16,000 *× g* for 5 minutes at 4°C. The supernatant was separated from the pellet and transferred into a new 1.5 mL tube with 40 μL glacial acetic acid, vortexed, and immediately transferred to Waters Sep-Pak tC18 columns conditioned with 6mL methanol and 6 mL water. Samples were allowed to flow through, and the columns were then washed with 3 mL hexanes and 6 mL water. Oxylipins were then eluted with 6mL methyl formate and 3 mL methanol, dried under nitrogen gas, and resuspended in 100 μL 1:1 MeOH:H_2_O. Samples were frozen at -80°C until analysis with mass spectrometry.

### Oxylipin LC/MS analysis

Extracts were separated on an Agilent 1290 Infinity II LC system or Bio LC System using either an Agilent ZORBAX Eclipse Plus C18, 150 × 2.1 mm, 1.8 μm column (Agilent 959759-902) or an Agilent Altura Eclipse Plus C18, 150 × 2.1 mm, 1.8 μm column (Agilent 204215-308) maintained at 40°C. No guard column was used for any setup. All instrumentation was maintained in temperature- and humidity-controlled environments at all times to limit day-to-day variability. All instrumentation was powered by uninterruptible power supplies. No columns were passivated. The chromatography gradient included mobile phase A containing 0.1% acetic acid in water, and mobile phase B containing 9:1 acetonitrile:IPA. The mobile phase gradient began with 85% mobile phase A, decreased to 5% over 21 minutes, then increased to 85% by 23 min and held for the remainder of the method run. The flow rate was maintained at 0.35 mL/min. The UHPLC system was connected to an Agilent 6495D triple quadrupole (QqQ) MS dual AJS ESI mass spectrometer. This method was operated in negative mode, with the iFunnel in Fragile mode (RF amplitude 50 V peak-to-peak on both the high- and low-pressure funnels) unless otherwise noted; Standard mode used 100 V on both funnels. The gas temperature was kept at 290°C with a flow of 11 L/min. The nebulizer was at 35 PSI, sheath gas temperature at 350°C, and sheath gas flow at 11 L/min. The VCap voltage was set to 3500 V, and the nozzle voltage was set to 1000 V. A dMRM inclusion list was used to individually optimize fragmentation parameters. Cycle time ranged from 96–500 ms, dwell time ranged from 10–500 ms, and the maximum concurrent MRMs was 42, with 250 total MRMs. Retention times varied between compounds from 0.8–2 min. Collision energies varied between compounds and ranged from 10–37 V. Transitions for all standards were generated with Agilent Compound Optimizer. The injection volume was 10 µL.

### Cell Culture

THP-1 cells (ATCC #TIB-202) were cultured in RPMI 1640 medium with 10% fetal bovine serum (Thermo Scientific A5256701) and 0.2% Primocin (Fisher Scientific ANT-PM-2). Monocyte-like THP-1 cells were collected by centrifuging cells and media at 250 *x* g for 5 minutes, washing with 10 volumes of PBS, and centrifuging again at 250 × g for 5 minutes. Cells were frozen at -80°C until analysis. M1 macrophage-like THP-1 cells were generated by incubating THP-1 cells in 100 nM PMA (Sigma-Aldrich P1585-1MG) for 48 hours, followed by a 24-hour incubation in growth media, then a 24-hour incubation in growth medium supplemented with 20 ng/mL of lipopolysaccharide (Cell Signaling Technologies 14011S) and interferon gamma (R&D Systems 285-IF-025). Cells were collected by scraping into 20:80 methanol:water and frozen at -80°C until analysis. Standard RPMI 1640 media was used for 10 mM glucose conditions. For creation of 20 mM glucose media, Dextrose (Millipore Sigma RDD016) was added to RPMI 1640 media, then passed through a 0.2 μm filter before adding FBS and Primocin as described above.

### LC Passivation

The stainless steel LC system was first replumbed to bypass the degasser before passivation. First, the solvent lines were flushed with 32.5% nitric acid for 6 hours, then flushed with water overnight. The pH of waste output water was checked for consistency with input water. Inlet filter frits for all pump channels were then replaced. The degasser was reconnected before any analysis.

### Data processing

Raw data was collected in .d format and checked manually in Agilent MassHunter Qualitative Analysis 10.0. The data was then uploaded to Agilent MassHunter QQQ Quantitative Analysis 10.1 for quantitation. A signal-to-noise ratio of 5:1 was used for the LLOQ, as specified in multiple reports.^9^ Chromatographic peaks also used a cutoff of at least 10 points along the peak for quantification. After manual inspection, data filtration, and integration as needed, peak height for standards and compounds were exported to .csv files. The value of the internal standard blank (ISB) was subtracted from each sample for each compound, and compound peak heights were normalized to the appropriate standard’s peak height, corresponding to a known concentration, to obtain compound concentrations in ng/mL. Compound-standard pairings are included in **Supporting Information, Table S2.** GraphPad Prism ver. 10 was used for figure generation and data analysis, including t-tests and ANOVA. RStudio was used for Venn diagram and heatmap generation. All files were uploaded to MassIVE (Accession number MSV000102273). The lipidomics standard reporting checklist for the project can be accessed at https://doi.org/10.5281/zenodo.23110687.

## 3. Methods, Results, and Discussion

Plasma samples from a single donor with known fasting status, health status, collection timeline, anticoagulation status, and time of day of collection were used for the initial experimentation. This allowed method development from a single donor sample, which allowed us to assess sample variability within our experiments (**Figure 2**). Additional pooled plasma was used to verify these initial observations. Solid-phase extraction was utilized for oxylipin analysis as it offers superior recovery and sample cleanup compared to liquid-liquid extraction.^19^ An example TIC of human plasma is shown in **Figure 2c**. We elected to use 39 different deuterated internal standards (**Table 1**) as different classes of oxylipins ionize differently, as observable in **Figure 2d**. Not all standards are easily visualized at the same TIC magnitude. All internal standard species were added in equal concentrations. Oxylipin extracts of same-donor plasma were pooled to achieve identical sample injections across experimental conditions; different replicates represent injection replicates. Separation was achieved by reverse-phase chromatography using mobile phases validated by others in the field.^9^

**Figure 2:**
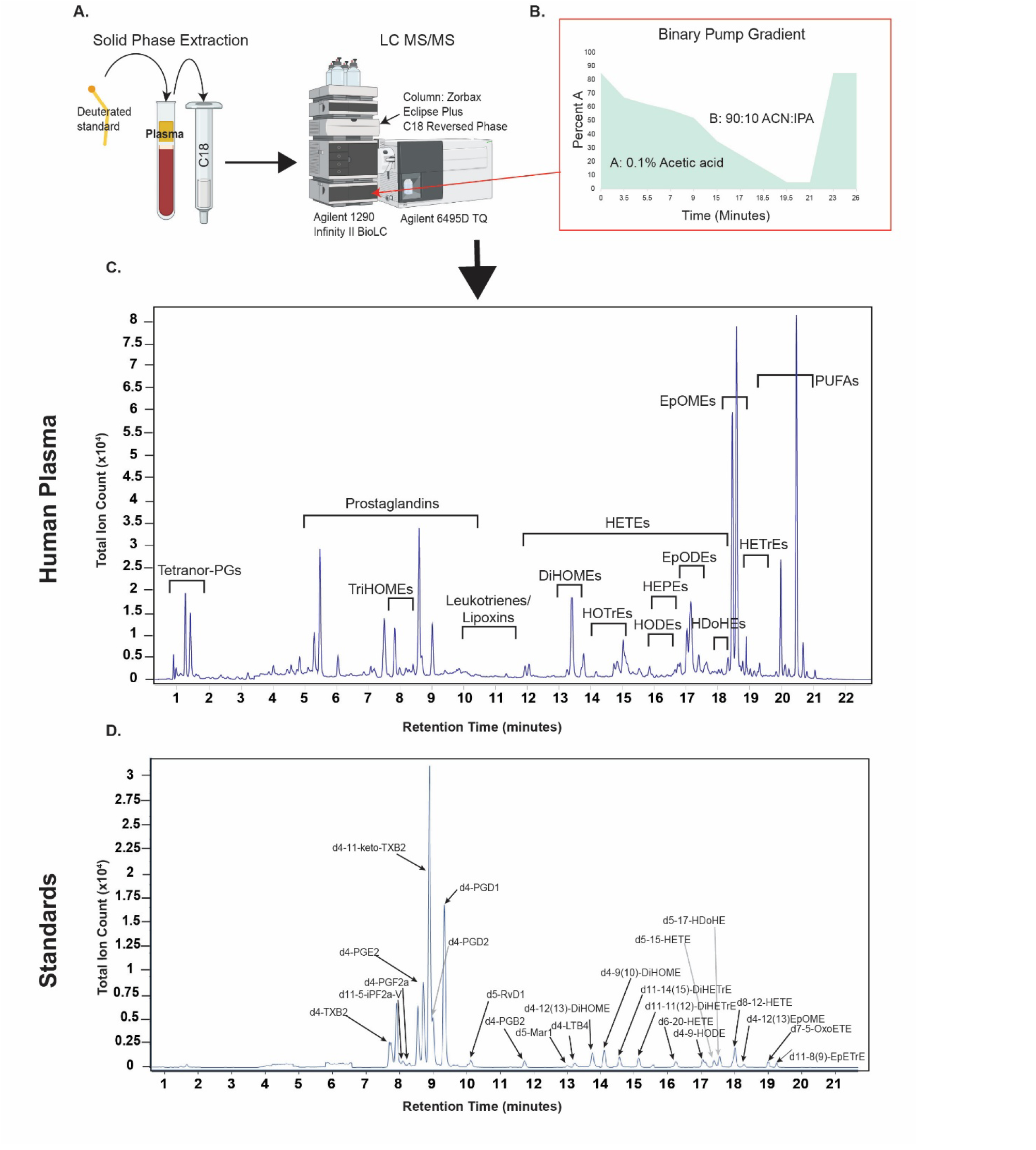
Oxylipin Extraction from Plasma and Mass Spectrometry Method. (A) Extraction method-deuterated oxylipin internal standards were added to plasma, and oxylipin samples were prepared with solid-phase extraction. (B) Binary pump gradient throughout the 26-minute run time of the method. (C) Total ion chromatogram of human plasma. (D) Total ion chromatogram of internal standards; not all standards are visualized at this TIC scale but are included in Table 1.

### 3.1 Impact of Ion Guide Voltages

Ion guide RF amplitude balances ion transmission against in-source fragmentation: higher voltages focus ions inside the optics of the instrument and can lead to fragmentation of analytes before reaching the first quadrupole.^16^ We therefore compared two ion funnel presets, the “Fragile” mode (RF voltage peak-to-peak = 50 V) and the “Standard” mode (100 V) (Figure 3a), using deuterated standards and a pooled plasma extract from a single donor. All comparisons used injection replicates (n = 3).

**Figure 3:**
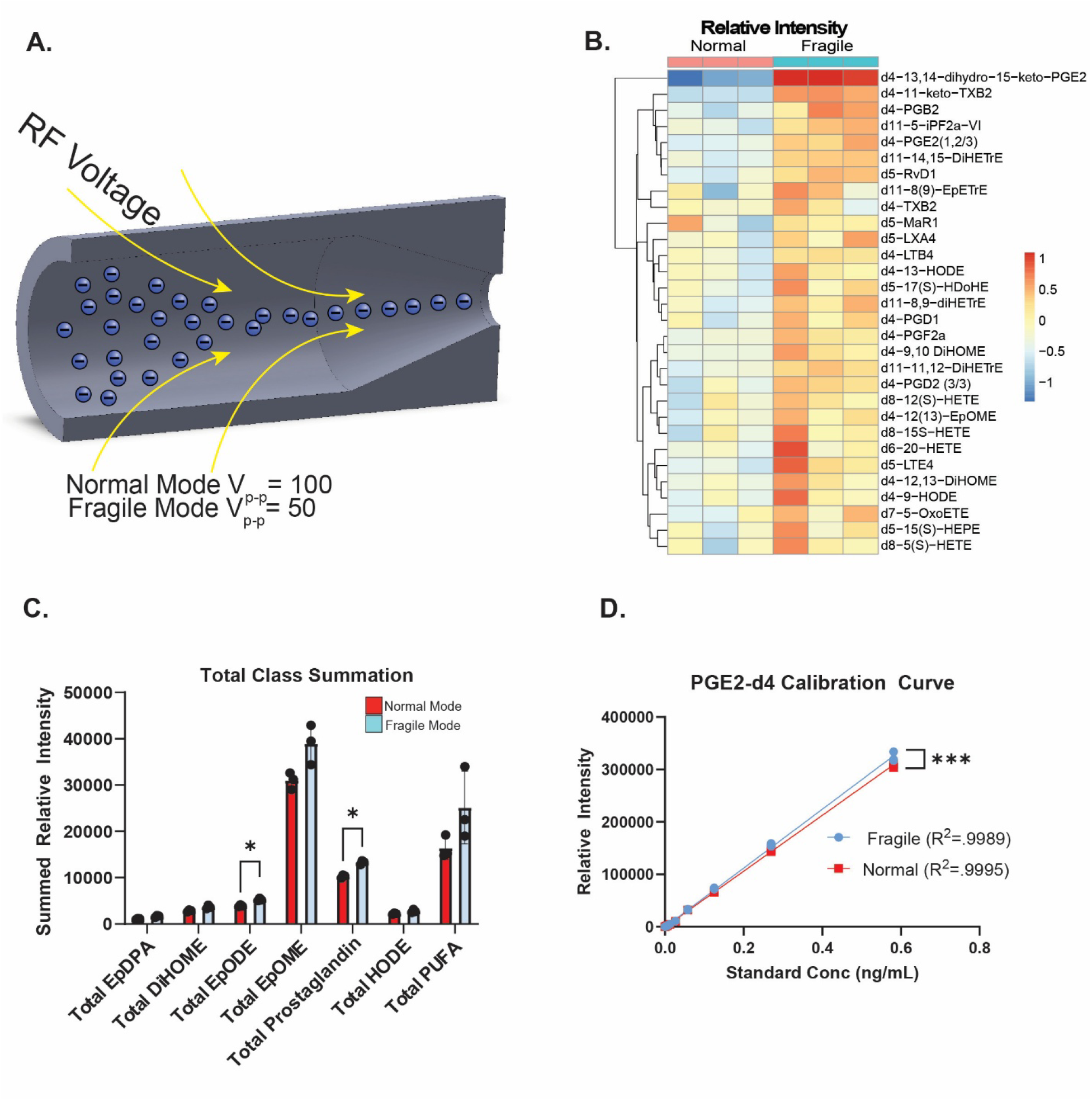
Lower Ion Funnel Voltages Produce Increases in Oxylipin Relative Intensities. Human plasma oxylipin extracts were run through the setup depicted in **Figure 2** with differing ion funnel voltages. (A) Ion funnel function visually represented, RF voltages applied for fragile and normal mode runs were 50V and 100V respectively. (B) Heatmap showing the top 25 most significantly changed oxylipin standard species between fragile and normal mode. (C) Total oxylipin class summations between normal (red) and fragile mode (blue), p-values calculated by two-tailed multiple unpaired t-test, n=3 injection replicates, *p<0.05. (D) Calibration curve showing injection concentration vs relative intensity of PGE2-d4 standard across normal mode vs fragile mode,***= p<0.001, Welch’s t-test.

We observed that fragile mode increased signal intensity in numerous oxylipin standards and endogenous oxylipins in plasma (**Figure 3b; Supporting Information, Figure S1a**). By class, epoxides and polyunsaturated fatty acids increased in relative intensity with lower ion funnel voltages, while most other classes had no change (**Figure 3c**). Peak shape, retention time, and peak quality were preserved across the species that changed significantly between fragile and normal mode (**Supporting Information, Figure S1d).** Calibration curves of deuterated standards spanning a range of concentrations were linear in both modes (**Figure 3d**), and fragile mode gave a higher response across multiple standards (**Supporting Information, Figure S1c**). This is a small but potentially meaningful change for the quantification of oxylipin species.

Hydroxylated fatty acids (hydroxyacids) were the only species with decreased intensity in fragile mode. This is consistent with manufacturer guidance that Fragile mode is optimized for labile analytes and low-m/z precursors, whereas Standard mode provides optimum transmission for most non-labile analytes,^16^ suggesting that hydroxyacids are comparatively resistant to in-source fragmentation. Hydroxyacids can be produced as intermediates of β-oxidation, lipid peroxidation of monounsaturated fatty acids, or possibly the breakdown products of certain oxylipins such as HETEs and HDoHEs.^20^ Hydroxylated MUFA- and PUFA-derived species are oxylipins by the definition used here, whereas saturated hydroxy fatty acids (e.g., 3OH-C05, 3OH-C12) and free fatty acids (e.g., C06, C09, C10) are related species; all are monitored because their abundance can indicate oxidation (see Section 3.3).

Using an additional pooled plasma sample from Innovative Research also showed increased oxylipin concentrations in fragile mode (**Supporting Information, Figure S1b**). Because concentrations are calculated relative to deuterated internal standards, we expected the standards and endogenous oxylipins to change in proportion, making this result unexpected. It could arise if deuterated oxylipin standards behave differently than endogenous oxylipins inside the ion guide, or if there is an inadequate number of standards and imperfect standard-compound pairing for relative analysis (Supporting Information, Table S2). Separately, absolute oxylipin levels in this commercial plasma may also be elevated by platelet-derived oxylipin formation during commercial plasma preparation and long-term storage (24 months at -80°C), which would not explain the difference between ion funnel modes.

Overall, these observations were consistent with previous literature that describes oxylipin stability and fragility as a difficult factor in analysis.^9^ A limitation of the instrument is that RF voltage is available only as presets, with no option to enter custom values that could be tuned for each compound of interest. Because most species were unaffected or improved by fragile mode, and switching between ion funnel modes lengthens cycle time, we opted to move forward with a method using only fragile mode for all species of interest in all subsequent experiments.

### 3.2 Impact of Bio-inert Surfaces

Next, we tested the effect of the bio-inert chromatography system on oxylipin quantification. The LC-MS system was used in the following configurations: Agilent Infinity II LC with Zorbax Eclipse plus C18 column (Standard_Steel), Agilent Infinity II Bio-inert LC with Zorbax Eclipse plus C18 column (Bio_Steel), and Agilent Infinity II Bio-inert LC with Altura Eclipse plus bio-inert coated C18 column (Bio_Altura). Sample processing, LC gradients, column temperature, autosampler temperature, and all other LC and MS parameters were held constant between the configurations. Setups containing the BioLC system, which is primarily composed of MP35N and titanium, were able to quantify new species, including hydroxylated fatty acid species, 8-HDoHE, 14(15)-EpETE, 19(20)-DiHDPA, and 11-HETE (**Figure 4a**).

**Figure 4:**
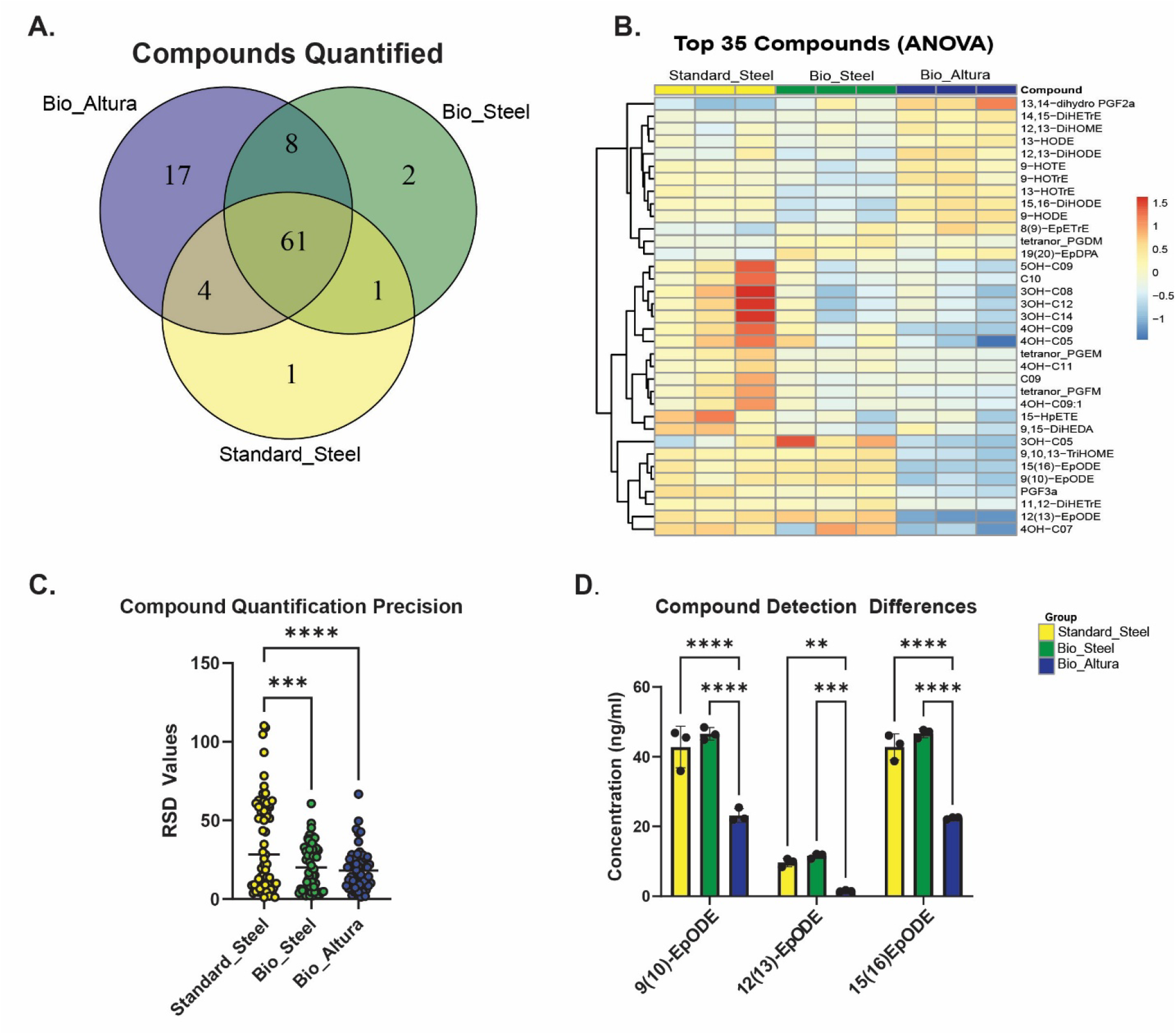
Fully Bio-Inert Chromatography Systems Improve Oxylipin Quantification and Precision. (A) Human plasma oxylipin extracts were run through 3 different systems: Standard_Steel (yellow, stainless-steel LC with stainless steel chromatography column), Bio_Steel (green, Bio-inert LC with stainless steel chromatography column), and Bio_Altura (purple, Bio-inert LC with Bio-inert Altura chromatography column). (B) Heatmap showing the top 35 most significantly different oxylipin species between chromatography setups. (C) Graph of relative standard deviation values for compounds detected in all three setups, p values calculated by Tukey’s multiple comparison testing, n=61, *p<0.05, **p<0.01, ***p<0.001, ****p<0.0001. (D) Bar graph showing EpODE species detected across the 3 chromatography setups, p values calculated by two-tailed multiple unpaired t-test, n=3 injection replicates, *p<0.05, **p<0.01, ***p<0.001, ****p<0.0001.

The Bio_Altura system resulted in more compounds surpassing the LLOQ than the other two, with compounds such as 12-HEPE, 12-HETE, PGE2, and LTD4 now quantifiable. Principal component analysis assessed the variance and spread of oxylipin quantification between the 3 chromatography systems. The Standard_Steel and Bio_Steel grouped together, while Bio_Altura was distinct (**Supporting Information, Figure S2a**). Heatmaps of compounds that were detectable in all systems showed significant differences in many compounds between systems (**Figure 4b**). Specifically, Bio_Altura detected higher levels of multiple species compared to stainless steel column-containing systems (**Figure 4b**), with higher levels of EpDPA, HODE, DiHOME, DiHODE, and HDoHE species. Conversely, Bio_Altura detected lower levels of PGF3α, 11,12-DiHETrE, and all EpODE species than those with stainless steel columns. Hydroxylated fatty acids were increased with the bio-inert chromatography column (**Supporting Information, Figure S2c**). The other pooled plasma dataset also showed more quantifiable oxylipins with the Bio_Altura and Bio_Steel systems (**Supporting Information, Figure S2b)** and increased concentrations for most species (**Supporting Information, Figure S2c).**

To assess precision in injection replicates between setups, we next plotted the relative standard deviation (RSD) values for only compounds detected in all three setups (**Figure 4c).** RSD values decreased for compounds in the Bio_Steel and Bio_Altura setups, suggesting that limiting stainless steel inside the LC stack can improve precision. However, many oxylipins had RSD values much higher than optimal for many compounds, even in the Bio_Altura setup. This is further evidence that oxylipin values must be interpreted with care and caution, as some compounds display high variation even in injection replicates. Of particular interest to us, EpODE species were significantly decreased with Bio_Altura (**Figure 4d**). To further evaluate our Bio_Altura setup, we performed calibration curves of several standards and observed high linearity in concentration vs relative intensity **(Supporting Information, Figure S2d)**. To ensure that quantification by height was not producing significant artifacts due to changes in peak shape, we quantified EpODE species with area and saw no significant difference compared to height **(Supporting Information, Figure S2e).** While previous reports have shown favorable effects on lipids, the few oxylipins quantified using different bio-inert setups were not significantly different from non-bio-inert setups.^13^ PEEK coatings versus other proprietary coatings were also shown to have different effects on lipid mass spectrometry.^13^ Our data support the use of bio-inert systems for oxylipin analysis.

### 3.3 Oxidation of PUFAs at Bio-Reactive Interfaces

The differences in epoxide concentrations detected among the stainless-steel and bio-inert setups were particularly intriguing and led us to consider the effect of stainless steel and iron on the generation of oxidized lipids within the LC system. Peroxidation of PUFAs, especially in the presence of iron, is a known and efficient reaction.^21–23^ This led us to test the effect of PUFA exposure to stainless steel on oxylipin levels detectable in mass spectrometry. Potentially, alpha-linolenic acid (ALA) could be converted into various EpODE and possibly other species within the chromatography instrument. To test this theory, physiologic concentrations of EpODE precursor (ALA, 20 μg/ml) alone were injected to quantify any potential epoxide formation.^24^ Both the Bio_Steel and Bio_Altura formed significant levels of EpODE species when only ALA was injected. Bio_Steel formed significantly more EpODEs, which was attenuated by adding 20 mg/L of butylated hydroxytoluene (BHT) into mobile phase B (**Figure 5a**).

**Figure 5:**
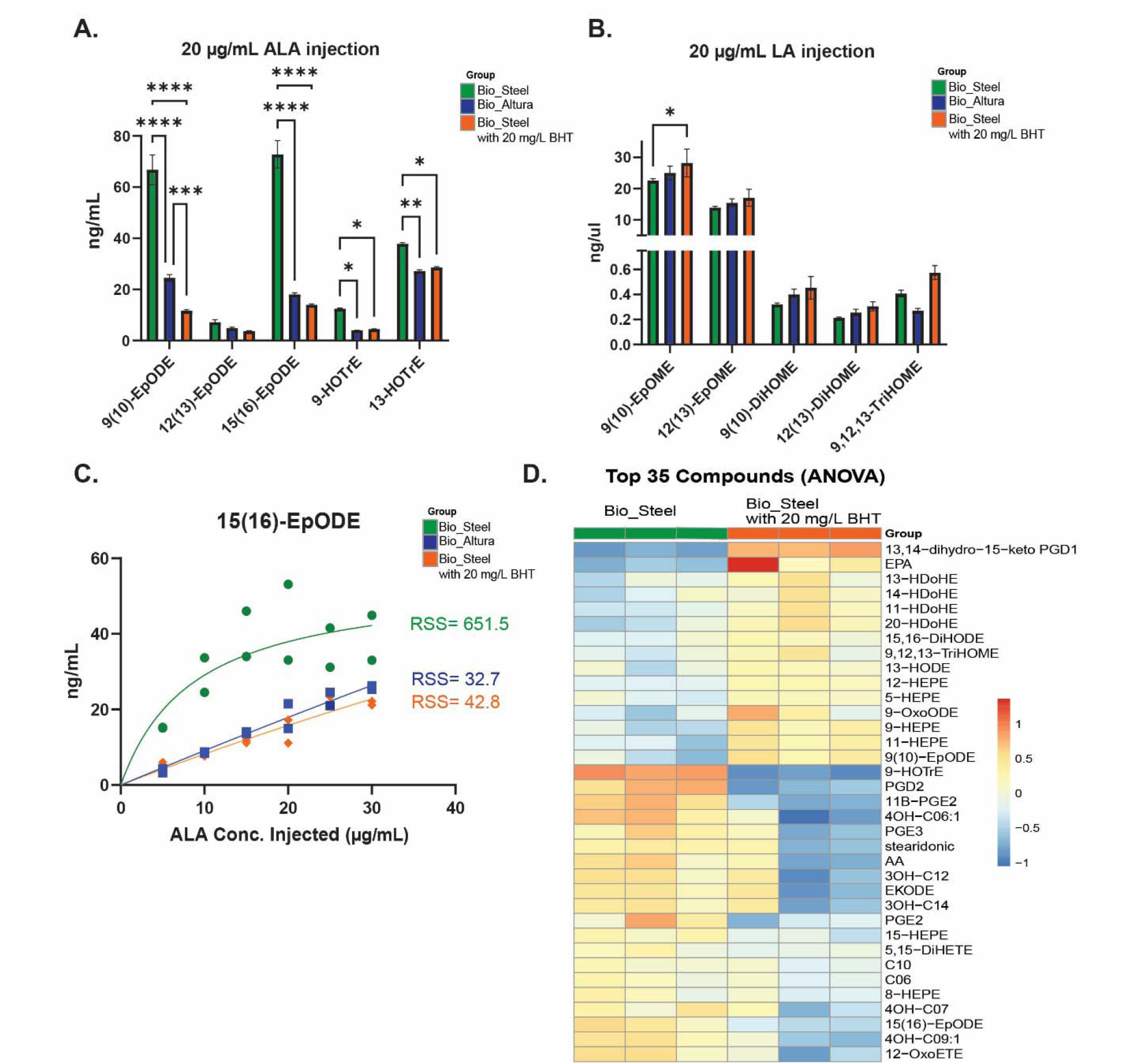
Stainless Steel Chromatography Columns May Modify Epoxide Oxylipins from Fatty Acids. (A) Bar graph comparing α-linolenic acid (ALA) injection into Bio_Steel (green), Bio_Altura (blue), and Bio_Steel with 20 mg/L BHT in mobile phase B (orange), assessing EpODE species and other ALA-derived oxylipins; p-values calculated by Tukey’s multiple comparison testing. (B) Bar graph comparing linoleic acid injection into Bio_Steel (green), Bio_Altura (blue), and Bio_Steel with 20 mg/L BHT in mobile phase B (orange), assessing EpOME species and other LA-derived oxylipins; p-values calculated by Tukey’s multiple comparison testing. (C) 15(16)-EpODE generation from ALA injection at various concentrations across the different chromatography setups. RSS= residual sum of squares for nonlinear lines of best fit. (D) Heatmap showing difference between human plasma analysis with Bio_Steel and Bio_Steel with 20 mg/L of BHT added to mobile phase B.

We hypothesized that adding BHT to the chromatography system would reduce any autooxidation reactions occurring, as BHT is a radical chain disruptor; this is the mechanism we propose for lipid oxidation within the LC.^18^ Controlling oxidation of samples before or during the extraction is the standard for quantifying physiologically relevant levels of oxylipins.^9,10,18^ BHT has been used in oxylipin analysis during the sample preparation step to limit autooxidation and the generation of oxylipin *ex vivo*, but recent reports have found this step unnecessary and even detrimental.^18^ One study even reported increasing the concentration of BHT by 10-fold but still observing autooxidation artifacts and *ex vivo* oxylipin formation.^18^ This suggests that the *ex vivo* formation could be occurring at steps beyond sample preparation, possibly inside the instrumentation itself. We also observed another class of ALA-derived oxylipins, HOTrEs, generated inside the instrumentation, an effect which was similarly limited by BHT addition and non-stainless steel column usage, providing further support for non-specific metal-catalyzed hydroperoxide modification of PUFAs (**Figure 5a**). This effect was minimized or reversed when utilizing linoleic acid (LA), another precursor of oxylipin synthesis (**Figure 5b**). Multiple classes of oxylipins arising from LA, including EpOMEs, DiHOMEs, and TriHOMEs were quantified at similar or higher levels when using bio-inert chromatography columns as compared to Bio_Steel or Bio_Steel with BHT. ALA is more susceptible to peroxidation than linoleic acid, as oxidizability increases with the number of bis-allylic methylene groups (CH2 groups between two double bonds). Therefore, in the linoleic acid injection, metals may be acting to further break down the baseline level of oxidized lipids and convert them away from detectable species rather than forming quantifiable epoxides or dihydroxy species, as LA is less readily oxidized through radical chemistry.^25^ Overall, it seems that the various PUFAs behave differently within each chromatography system. This formation of epoxides was confirmed to be precursor-specific **(Supporting Information, Figure S3a, S3b)**.

Next, we determined that EpODE formation from ALA is concentration-dependent (**Figure 5c**), with higher injected ALA concentrations producing more 15(16)-EpODE. Variability around the fitted dose–response curve, quantified as the residual sum of squares (RSS), was greater with the Bio_Steel setup than with Bio_Altura or Bio_Steel with BHT. This led us to hypothesize that the variation and decreased precision of Bio_Steel come from oxidation reactions, as the precision can be increased by adding BHT. It is reasonable to infer that detected differences in EpODE species in human samples could be due to differences in ALA levels. While this system lacks the ligand-complexed iron and peroxide oxidant used in synthetic epoxidation catalysis, metal-accelerated autoxidation of ALA could still occur, albeit at a lower rate. Previous work has shown that a variety of non-complexed metal ions, including iron, have the ability to oxidize lipids when samples are extracted and incubated with these ions before LC-MS analysis.^26^ This is another reason why stainless steel chromatography systems are often passivated, as stainless steel columns have been known to leach metal ions and oxidize or degrade amino acids.^27^ Significant differences in detected levels of oxylipin species with passivation of our Standard_Steel system were observed, especially a decrease in 15(16)-EpODE **(Supporting Information, Figure S3d).** EpODE species were still generated inside the fully bio-inert setup. Because the Bio LC flow path is primarily MP35N (nickel, cobalt, chromium, and molybdenum) and titanium, and “bio-inert” refers to reduced analyte adsorption rather than redox inertness, this may represent radical peroxidation catalyzed by other transition metals. Alternatively, analytical grade ALA may also contain a significant level of EpODEs that are detectable with high-sensitivity mass spectrometry, so detecting some level of EpODEs is not unexpected.^28^ Because EpODEs formed at similar levels in the Bio_Steel and Bio_Altura setups, a metal-dependent background shared by both setups would not have been apparent in our initial comparison of the setups.

Finally, we added BHT to mobile phase B as done in **Figure 5a** and analyzed plasma samples. We observed differences in quantification in the single-donor plasma (**Figure 5d; Supporting Information, Figure S3c).** Notably, the decrease in EpODE species observed with Bio_Altura and BHT addition was recapitulated, with 15(16)-EpODE showing decreases in both plasma datasets. This provides further support for spontaneous metal-catalyzed hydroperoxide oxylipin formation within the LC system, and it is likely not just limited to EpODE species, as many other oxylipin species showed changes when BHT was added. We also observed a decrease in formation of 4-hydroxy fatty acids, leading us to postulate that certain hydroxy fatty acids can be formed by the same peroxide mechanism inside the system. This finding is consistent with our theory of fatty acids being peroxidized or significantly affected/converted by the conditions and metals exposed in chromatography systems. Additionally, when ALA was injected through LC instruments without chromatography columns to assess EpODE generation, there was less 15(16)-EpODE with the bio-inert LC as compared to standard LC setup, providing further evidence that exposed stainless steel contributes to formation (**Supporting Information, Figure S3e)**. To verify that we were detecting 15(16) EpODE, we compared the mass spectrum of our 15(16)-EpODE to a previously generated 15(16)-EpODE mass spectrum and observed high overlap **(Supporting Information, Figure S3f)**, helping to strengthen our theory of 15(16) EpODE generation.^29^ The Bio_Steel system continued to display decreased precision in repeated injections which was remedied with Bio_Altura. We remained unsure of the biological significance or accuracy of adding BHT to mobile phases, as BHT has the potential to increase ion suppression effects or adduct formation. Therefore, we elected to continue using Bio_Altura and standard mobile phases going forward.

### 3.4 Application Across Human Biological Matrices

To test this method’s ability to isolate oxylipins from other human tissues, we employed blood fractionation to separate individual components of whole human blood. Granulocytes, peripheral blood mononuclear cells, and erythrocytes were separated, and oxylipins were analyzed as above. Overall, the method was able to detect 56 oxylipin species in the different blood fractions. 25 species were shared between all the cell types, and 31 species varied between the different cell types, with specific cell types displaying unique oxylipins (**Figure 6a**). The erythrocytes displayed small but present levels of many oxylipins that overlapped with the rest of the blood fractions. This could represent platelet or granulocyte contamination of the RBC fraction. Platelets are a common contaminant of Ficoll-isolated fractions and the main source of 12-lipoxygenase products (12-HETE, 12-HEPE, and 12-oxoETE); because platelet contamination was not assessed (e.g., by CD41/CD61 staining or platelet counts), platelet-derived oxylipins cannot be excluded. Red blood cell lysis also releases heme, which efficiently catalyzes the decomposition of fatty acid hydroperoxides,^22^ and no antioxidant was added during fractionation. Ex vivo formation is therefore still very possible, as Ficoll-Paque fractionation requires 30 minutes of room-temperature exposure at each of several steps, so the erythrocyte and granulocyte profiles should be interpreted with caution. ^30,31^

**Figure 6:**
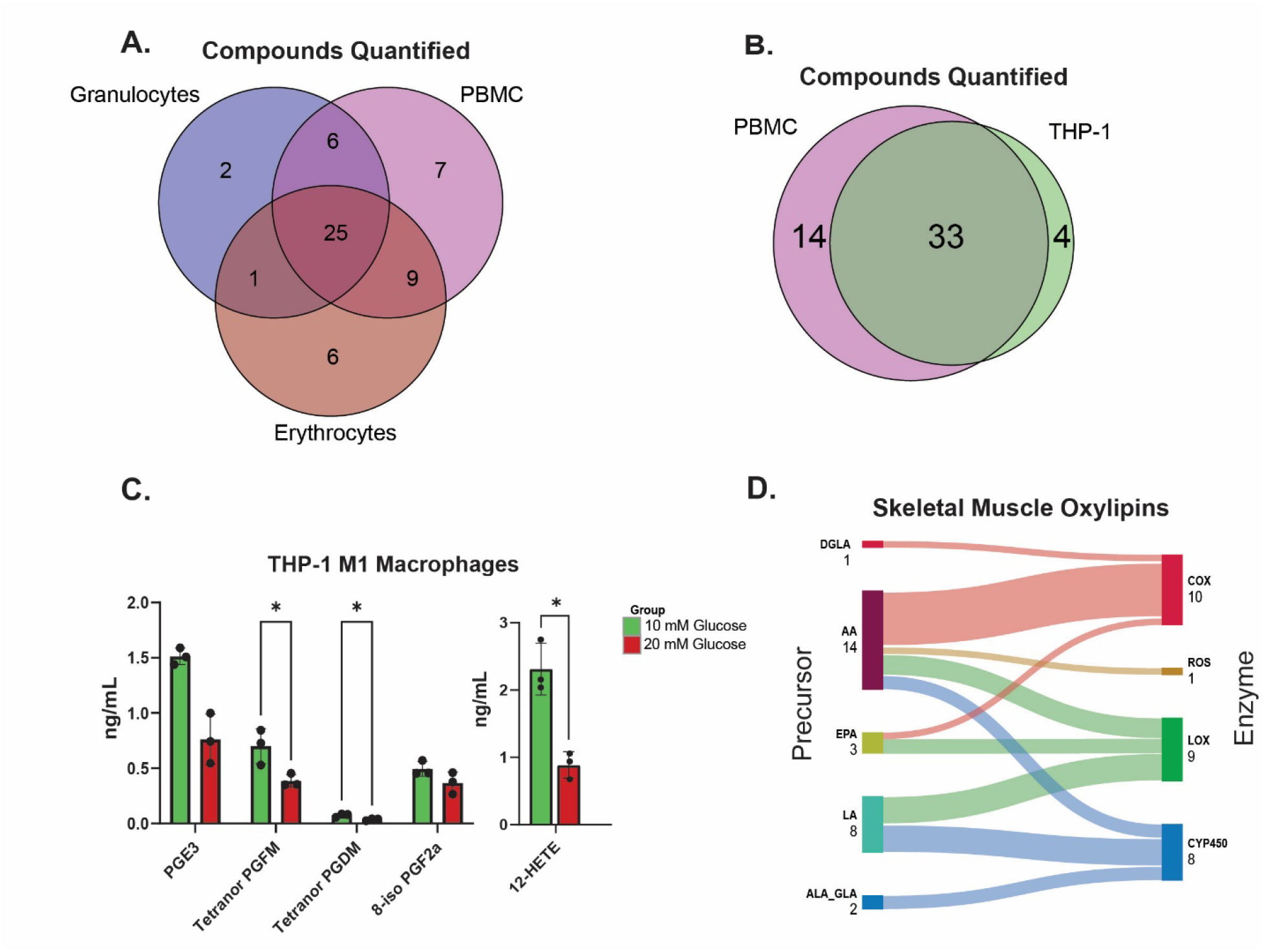
Application of Oxylipin Method Across Human Biological Matrices. Bio_Altura assessment between sample matrices. (A) Venn diagram of human blood fractions of granulocytes (blue), peripheral blood mononuclear cells (PBMC, purple), and erythrocytes (orange). (B) Venn diagram of PBMCs (purple) and THP-1 cells (green), a human-derived monocyte line. (C) Oxylipin profiles of hyperglycemic vs normoglycemic M1 macrophages differentiated from THP-1 cells, n=3, p-values calculated by two-tailed multiple unpaired t-test. (D) Sankey plot showing oxylipins quantified as categorized by precursor fatty acid and enzyme of synthesis

To further investigate the Bio_Altura method with blood fractions, we studied standard separation in plasma fractionation, extraction recovery, and matrix effects in our setup. Standards were spiked into whole blood, and then blood was fractionated using the Ficoll-Paque extraction described above. We observed that the plasma fraction contained most of the standards spiked into whole blood, showing that blood fractionation is effective at sequestering extracellular oxylipins into the plasma fraction, but a small amount of standards may bind or become internalized in cells **(Supporting Information, Figure S4a)**. Next, we tested the extraction efficiency of our method by comparing standards spiked into our system vs. an equal amount of standards that were taken through the extraction using plasma matrix for both groups. We observed a recovery level for most standards around 55%, which is lower than other lipidomics extractions, but possibly consistent with previous reports using oxylipin solid-phase extraction **(Supporting Information, Figure S4b)**.^19^ Finally, we tested the effects of matrix on standard relative intensity by spiking in standards to plasma matrix or methanol. We observed ion enhancement for most classes of standards, consistent with previous literature **(Supporting Information, Figure S4c).**

We also evaluated CUDA (12-(3-cyclohexylureido)dodecanoic acid) as an extraction additive to determine whether soluble epoxide hydrolase inhibition during sample handling altered epoxide and diol oxylipin measurements. CUDA-containing extractions showed similar overall oxylipin profiles and technical reproducibility compared with the standard BHT-containing extraction condition, but did not broadly increase epoxide signals or decrease downstream diol signals **(Supporting Information, Figure S4d)**. Therefore, CUDA was not incorporated into the final extraction method. Technical precision was assessed using plasma extracts. Across triplicate measurements, 70 analytes were detected, with a median CV of 4.1%; 67 of 70 detected analytes had CV values ≤20% **(Supporting Information, Figure S4e).** We also evaluated short-term autosampler injection stability by reinjecting plasma extracts at 6 h and 12 h. Median responses were 98.9% and 94.2% of the initial plasma mean at 6 h and 12 h, respectively, indicating good short-term stability **(Supporting Information, Figure S4f).** Inter-day precision was evaluated using pooled plasma QC samples analyzed in triplicate on three separate days, yielding a median between-day CV of 12.8%, with 73.4% of analytes meeting the ≤20% CV criterion **(Supporting Information, Figure S4e).** These results demonstrate encouraging reproducibility across analytical days.

Next, we compared the oxylipins found in a human-derived monocyte line, THP-1, to peripheral blood mononuclear cells (PBMCs) found in human samples. We observed high overlap in the species detected in the PBMCs and THP-1 monocytes, with 33 of the 51 detected species being identified in both cell populations (**Figure 6b).** Cultured cells have limited availability of oxylipin precursors, specifically PUFAs, because of their low concentrations in the standard FBS used in cell culture, so observing highly similar oxylipin profiles with fewer species detected in THP-1 cells was consistent. We then compared THP-1–derived M1 macrophages cultured in hyperglycemic (20 mM glucose) vs normoglycemic (10 mM glucose) conditions to determine this method’s capacity to uncover treatment effects. THP-1 monocytes have been shown to differentially regulate COX enzymes in response to hyperglycemia, but specific oxylipin differences have not yet been shown.^32^ We observed differences across various prostaglandin species. Specifically, PGE3, tetranor PGFM, and tetranor PGDM were significantly decreased in hyperglycemic conditions relative to normoglycemic conditions (**Figure 6c**). 12-HETE also displayed decreases in the hyperglycemic THP-1 cells. We expected to see differences in prostaglandins, as COX-2 is a key enzyme that increases in expression and protein levels in response to hyperglycemia.^32^ However, the direction of change surprised us, which was possibly due to these cells being differentiated into M1 macrophages.

Finally, we analyzed skeletal muscle tissue for oxylipins and observed fewer detectable species as compared to other tissues, although oxylipins synthesized from a variety of precursors and enzymes were detectable (**Figure 6d).** We quantified 18 different hydroxylated fatty acids from muscle tissue. We also obtained small quantities of human beta-islet tissue to test our method as well. A full list of all quantified compounds, including those from human beta-islet tissue, is shown in **Supporting Information, Table S1.** Two specialized pro-resolving mediators (SPMs), RvD4 and RvD5, were quantified in muscle and RBC samples, respectively. This was unexpected and may have been a result of sample autooxidation, especially as RBCs contain large amounts of iron. Because ex vivo and in-instrument oxidation can generate SPM-like signals, these two assignments should be interpreted with caution. Chromatographic peaks annotated with signal-to-noise, retention time, collision energies, and MS transitions for both SPMs and standards are included in **Supporting Information, Figure S5.** We also analyzed the ability of our chromatography system to separate critical oxylipin pairs and classes by following guidelines published previously.^9^ Overall, we saw adequate separation of DiHODE, OxoETE, HpETE, HDoHE, and PG oxylipin classes **(Supporting Information, Figure S6).** Previous studies analyzing oxylipin profiles of human blood-derived cells reported little to no pro-resolving oxylipins in PBMCs, granulocytes, or whole blood, which differs slightly from our results.^33^ Observing fewer low-abundance pro-resolving mediators may have been due to several factors in other studies, such as differences in the anticoagulant used during blood collection (heparin vs EDTA), stainless steel column usage, variability due to normal human physiology, and different instrumentation.^14^ Using human samples and human-derived cell lines, we were able to use this method to quantify oxylipins in a variety of tissues and show a treatment effect in human cell lines that is consistent with previous literature.

## 4. Conclusions

We first demonstrated that decreases in ion funnel voltage improve relative intensity of detected oxylipins while preserving peak shape. We also show that usage of fully bio-inert setups with bio-inert chromatography columns improves both detection and precision of oxylipin quantification. Furthermore, we observed that utilizing stainless steel columns in chromatography systems modifies the levels of oxylipins quantified, specifically EpODE species, possibly through metal-catalyzed oxidation reactions. We recommend limiting oxidation reactions when analyzing oxylipins through the use of bio-inert systems and columns. More work is needed to assess if adding antioxidants, such as BHT, directly into chromatography mobile phases is a viable solution for labs without access to bio-inert systems. Finally, we demonstrate this method’s utility in quantifying oxylipins from a variety of human tissues, including erythrocytes, granulocytes, peripheral blood mononuclear cells, muscle, and THP-1 cells. Using this new system, we hope to discover more species of oxylipins that are physiologically accurate and take advantage of the increased precision to identify species that correlate with or prognosticate disease. These results demonstrate that oxylipin analysis has improved precision and compound detection with bio-inert chromatography. To achieve the most physiologically accurate oxylipin analysis with LC-MS, oxylipin quantification should use bio-inert LC systems; if this is unattainable, in-instrument oxidation should be assessed.

## Supporting Information

Supporting Information (PDF): Figures S1–S6 (ion guide parameters; bio-inert chromatography setups; PUFA oxidation in stainless-steel-containing setups; Bio_Altura method evaluation; chromatograms of detected specialized pro-resolving mediators; chromatographic separation of critical oxylipin pairs) and Tables S1–S2 (compounds quantified in six human tissues; compound–standard pairings).

## Author Contributions

J.T.S., L.W., J.M., and J.A.S. conceptualization; J.T.S., L.W., and J.M. methodology; J.T.S. and L.W. data analysis; all authors investigation; J.T.S., J.M., and J.A.S. writing and editing.

## Conflict of Interest

F.T. is affiliated with Agilent Technologies, Inc., the manufacturer of the instrumentation, columns, and software presets evaluated in this study. The remaining authors declare no competing interests.

## Acknowledgments

We would like to thank Dr. Craig Wheelock for his advice on internal standards and methods. We thank all Simcox Lab members for assistance with experiments, editing materials, and thoughtful discussions. Dr. Simcox is an HHMI Freeman Hrabowski Scholar. Research reported in this publication was supported by the University of Wisconsin-Madison Physician Training Grant (T32GM140935 to JTS and MM); William H. Peterson Fellowship in Biochemistry (to JTS); Breakthrough T1D (JDRF201309442 to JAS); R01 through NIH/NIDDK (R01DK133479 to JAS); R01 through NIDDK (R01DK137976 to JAS); NIH/NIA (U01AG076941 to ARK); R01 through NIDDK (R01DK137505 to MEK); Merit Review Award through the Department of Veterans Affairs (I01BX005804); Research Career Scientist Award through the Department of Veterans Affairs (IK6BX006804); and Wisconsin Nathan Shock Center of Excellence in the Basic Biology of Aging (P30AG092586). The content is solely the responsibility of the authors and does not necessarily represent the official views of the National Institutes of Health. The work was also supported in part by startup funds from the University of Wisconsin-Madison School Department of Biochemistry to JAS.

## Abbreviation

AJS: Agilent Jet Stream
ALA: alpha-linolenic acid
BHT: butylated hydroxytoluene
COX: cyclooxygenase 12-(3-cyclohexylureido)dodecanoic
CUDA: acid
CV: coefficient of variation
CYP450: cytochrome P450
DiHDPA: dihydroxydocosapentaenoic acid
DiHETrE: dihydroxyeicosatrienoic acid
DiHODE: dihydroxyoctadecadienoic acid
DiHOME: dihydroxyoctadecenoic acid
dMRM: dynamic multiple reaction monitoring
EDTA: ethylenediaminetetraacetic acid
EpDPA: epoxydocosapentaenoic acid
EpETE: epoxyeicosatetraenoic acid
EpODE: epoxyoctadecadienoic acid
EpOME: epoxyoctadecenoic acid
ESI: electrospray ionization
FBS: fetal bovine serum
HDoHE: hydroxydocosahexaenoic acid
HEPE: hydroxyeicosapentaenoic acid
HETE: hydroxyeicosatetraenoic acid
HODE: hydroxyoctadecadienoic acid
HOTrE: hydroxyoctadecatrienoic acid
HpETE: hydroperoxyeicosatetraenoic acid high performance liquid
HPLC: chromatography
IPA: isopropyl alcohol
ISB: internal standard blank
LA: linoleic acid liquid chromatography-mass
LC-MS: spectrometry
LLOQ: lower limit of quantification
LOX: lipoxygenase
LTD: leukotriene
MAPK: mitogen-activated protein kinase
MeOH: methanol
MRM: multiple reaction monitoring
MUFA: monounsaturated fatty acid
OxoETE: oxoeicosatetraenoic acid
PBMC: peripheral blood mononuclear cell
PBS: phosphate buffered saline
PEEK: polyether ether ketone
PFAS: per- and polyfluoroalkyl substances
PG: prostaglandin
PGDM: prostaglandin D2 metabolite
PGE: prostaglandin E
PGF: prostaglandin F
PGFM: prostaglandin F2 alpha metabolite
PMA: phorbol 12-myristate 13-acetate
PUFA: polyunsaturated fatty acid
QC: quality control
RBC: red blood cell
RF: radiofrequency
RSD: relative standard deviation
RSS: residual sum of squares
RvD: resolvin D
SPE: solid phase extraction
SPM: specialized pro-resolving mediator
TIC: total ion chromatogram
TriHOME: trihydroxyoctadecenoic acid ultra high performance liquid
UHPLC: chromatography

## For Table of Contents Use Only

**Brief Synopsis:** Bio-inert (MP35N) liquid chromatography columns limit the oxidation of α-linolenic acid and other polyunsaturated fatty acids, which stainless-steel-containing LC systems can convert into oxylipins; lower ion funnel RF voltage further improves oxylipin detection.

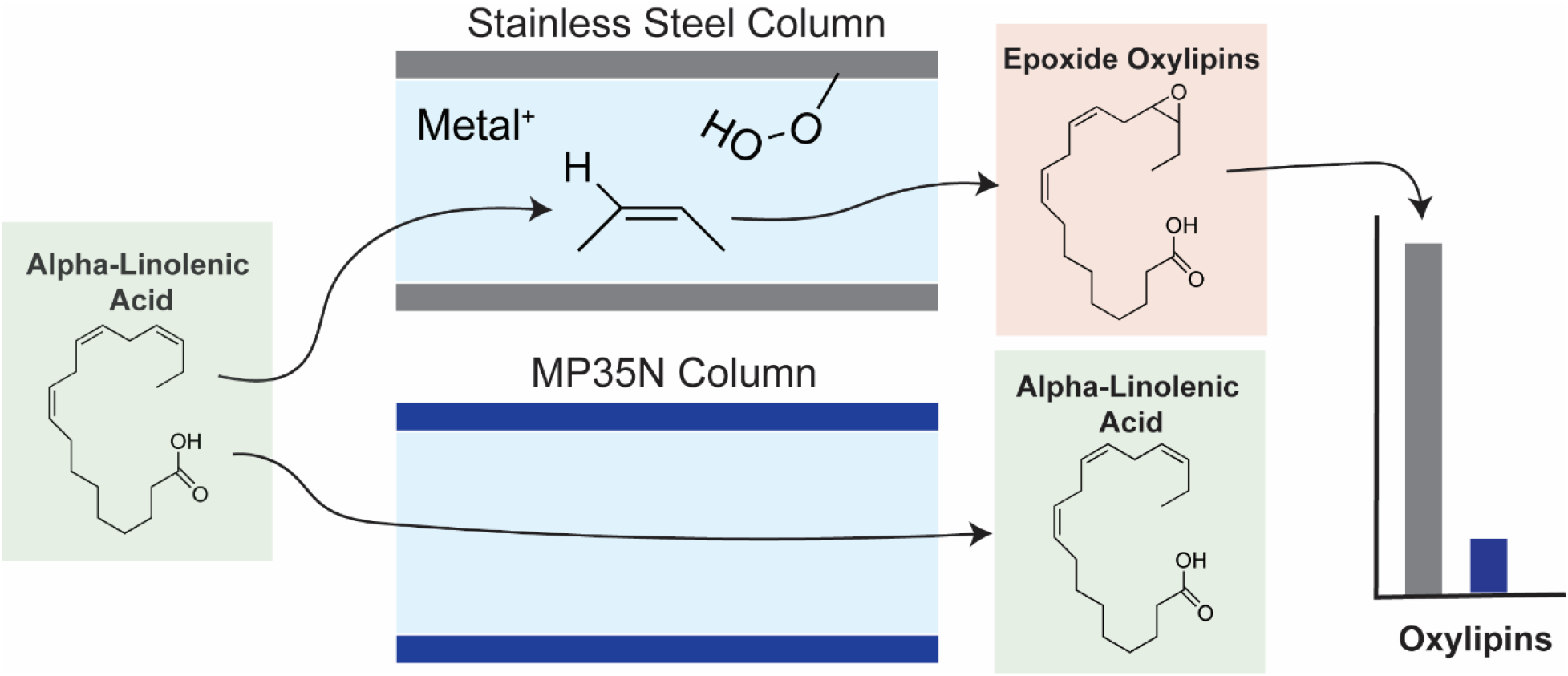

## Notes

https://explorer.gnps2.org/MSV000102273?dataset_accession=MSV000102273&metadata_source=MASSIVE&metadata_option=

### Competing Interest Statement

The authors have declared no competing interest.

### Summary of Updates

We have strengthened the manuscript through several major changes, including: New Figure 1: Schematic of oxylipins; added figure panels for main Figures: Figure 2C, Figure 3D, Figure 4B & C, Figure 5 B & D; Added Figure S4 for suggested method validation; Added Figure S5 for chromatograms of all detected SPMs; Added Figure S6 for chromatographic separation of oxylipins; and Added Table S2 for compound-standard pairings.

