## Supplemental information for "Optimizing Oxylipin Analysis with Liquid Chromatography Mass Spectrometry through Bio-Inert Systems and Ion Funnel Adjustments"

oxidation, HPLC, inflammation, human

**Corresponding author**

*

| **Title** | **Page Number** | **Main Figure of Support** | **Title** |
| --- | --- | --- | --- |
| Figure S1 | S3 | 3 | Ion guide parameters alter oxylipin detection |
| Figure S2 | S5 | 4 | Fully bio-inert chromatography setups improve oxylipin quantification |
| Figure S3 | S7 | 5 | PUFAs may be modified through oxidation in stainless-steel containing chromatography setups |
| Figure S4 | S9 | 6 | Bio_Altura method investigation |
| Table S1 | S11 | 6 | Lists of compounds quantified using oxylipin analysis in 6 different human tissues |
| Figure S5 | S13 | 6 | Chromatograms of all detected SPMs |
| Figure S6 | S14 | 6 | Separation ability of the chromatography system for oxylipin analysis. |
| Table S2 | S15-S16 | 6 | Compound-standard pairings for oxylipin analysis |

**Supporting Results and Discussion**

Human plasma was used to evaluate normal (RF voltage of ion guides at 100V peak-peak) and fragile (50V peak-peak). Overall, fragile mode increased compound relative intensities, as seen in **Figure S1a.** Using pooled plasma, we also observed that compound ng/mL quantified using relative quantification were increased in some species **(Figure S1b)**. This effect may have been the result of deuterated oxylipin standards behaving differently than endogenous oxylipins inside of ion guide machinery. This may also be the result of an inadequate standard representation and imperfect standard-compound pairing for relative analysis. Additional calibration curves for standards obtained using normal and fragile modes are visible in **Figure S1c.** All compounds achieved adequate linearity, as assessed through R^2^ coefficients, and 12-HETE-d8 and RvD1-d4 displayed increased relative intensities using fragile mode, as assessed by Welch’s t-test for difference in slope between lines. We also observed that peak shape was maintained and increased in height for certain compounds when using fragile mode **(Figure S1d).** This expands on **Figure 3** in the main manuscript.

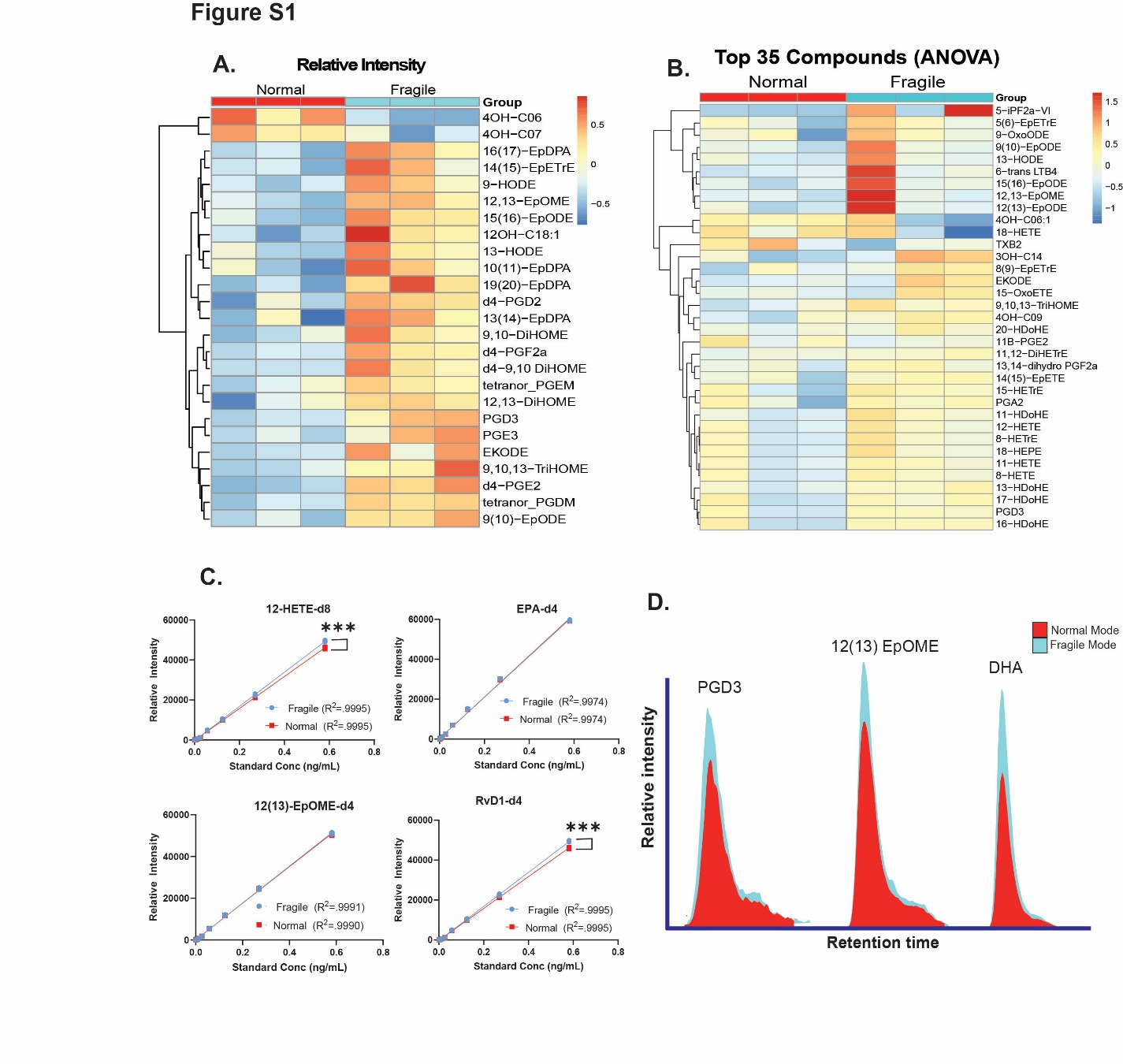

**Figure S1:** Ion guide parameters alter oxylipin detection. (A) Relative intensities for the 25 most significantly changed oxylipins in normal vs. fragile ion guide modes in human plasma. (B) Pooled plasma compound levels as assessed through relative quantification in normal vs fragile mode. (C) Calibration curves of different oxylipin standards in different ion guide modes, ***= p<0.001, Welch’s t-test. (D) Chromatographic peaks for representative samples showing peak shape as related to ion guide mode.

Next, we made different chromatography setups Standard_Steel (stainless-steel LC with stainless steel chromatography column), Bio_Steel (Bio-inert LC with stainless steel chromatography column), and Bio-Altura (Bio-inert LC with Bio-inert Altura chromatography column) to test the effects of LC setup on oxylipin analysis. We utilized principal component analysis to analyze the differences between the 3 chromatography systems based on the oxylipin levels quantified in all 3 setups **(Figure S2a)**. We observed that Standard_Steel and Bio_Steel grouped, while Bio_Altura was separated from the other two setups. Next, we analyzed a pooled plasma sample obtained from Innovative Research for secondary validation to ensure the effects observed were not unique to only our plasma. As seen in our healthy donor plasma, we observed increased species quantifiable when using the Bio_Altura setup as compared to Bio_Steel, showing that the bio-inert chromatography column was able to quantify more compounds as seen in our healthy donor plasma **(Figure S2b)**. Also similar to our healthy donor plasma, we observed that compounds were quantified in different levels using the bio-inert chromatography column **(Figure S2c).** To validate that Bio_Altura was suitable for compound detection across a variety of concentrations, we ran calibration curves of oxylipin standards across multiple classes and observed good linearity in injected concentration vs. relative intensity across a range of concentrations **(Figure S2d).** Finally, to verify that quantifying by height did not produce any confounding effects due to differing peak shape between setups, we quantified EpODE species by area and compared it to height. We observed no difference, showing that height quantification may be noninferior to area quantification when comparing chromatography setups. This expands on **Figure 4** in the main manuscript.

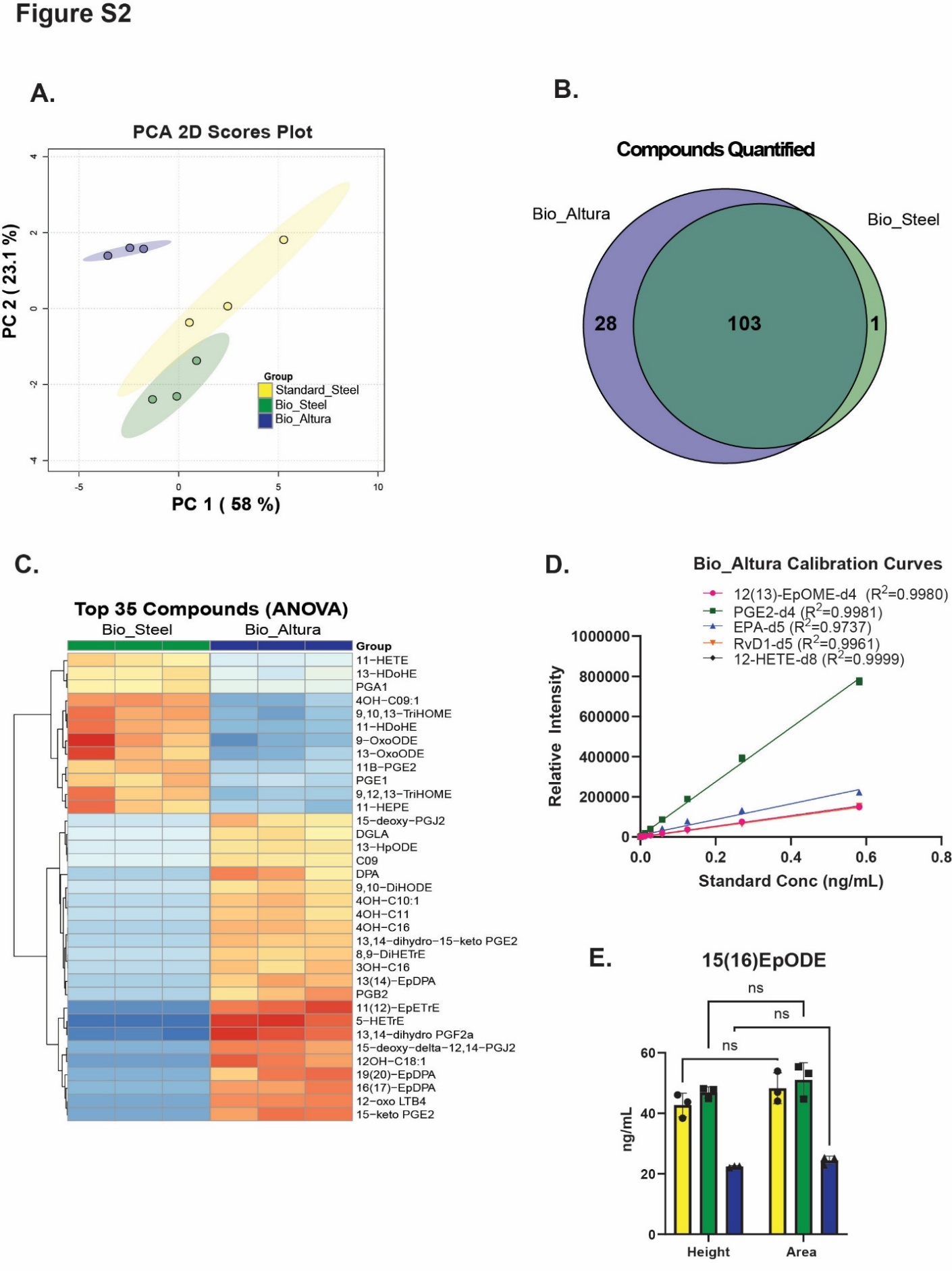

**Figure S2** Fully bio-inert chromatography setups improve oxylipin quantification. (A)Principal component analysis for the three setups Standard_Steel (yellow, stainless-steel LC with stainless steel chromatography column), Bio_Steel (green, Bio-inert LC with stainless steel chromatography column), and Bio-Altura (purple, Bio-inert LC with Bio-inert Altura chromatography column) for compounds quantified in all 3 setups only. (B) Compounds quantified in Bio_Altura vs Bio_Steel using pooled plasma as a secondary validation. (C) Heatmap of the top 35 significantly different compounds quantified in Bio_Altura vs Bio_Steel in pooled plasma. (D) Calibration curves of select oxylipin standards using Bio_Altura. (E) 15(16)-EpODE quantified using area vs height peak integration in various chromatography setups (Tukey’s multiple comparison testing).

To study oxidation in our chromatography system, we injected analytical-grade standards of various PUFAs into our chromatography setups. We observed virtually no epoxides being formed inside the instrument when their precursor was not injected **(Figure S3a, S3b).** When ALA was injected, no LA-derived epoxides were formed, and vice versa, showing the epoxide formation was specific to the precursor injected and not a random MS artifact generated from instrumentation. To further study the effect of oxidation on plasma analysis, we used the pooled plasma to test whether BHT added to the chromatography system would change detection of oxylipins in human plasma. The healthy donor plasma showed significantly different levels of multiple oxylipins when BHT was added to the Bio_Steel setup (**Figure S3c),** implying that controlling oxidation may have significant effects on oxylipin analysis**.** We then passivated our stainless steel chromatography system and compared the effects of passivation on oxylipin quantification **(Figure S3d).** Additionally, we injected ALA through LC instruments without chromatography columns to determine if we could observe an effect on EpODE generation without columns (**Figure S3e).** We observed less 15(16)-EpODE generation with the bio-inert LC as compared to standard LC setup, providing further evidence that exposed stainless steel modifies levels of certain oxylipins. Passivation and the creation of a less-bioactive layer on the surface of the stainless steel significantly affected the levels of detected oxylipins. To validate that the species we generated inside the mass spectrometer was 15(16) EpODE, we again injected ALA into the Bio_Steel setup and generated a mass spectrum of the species that was generated. It had high similarity to previous spectra of 15(16) EpODE, allowing us to strengthen the theory that we did generate 15(16)-EpODE inside our LC instrument. This expands on **Figure 5** in the main manuscript.

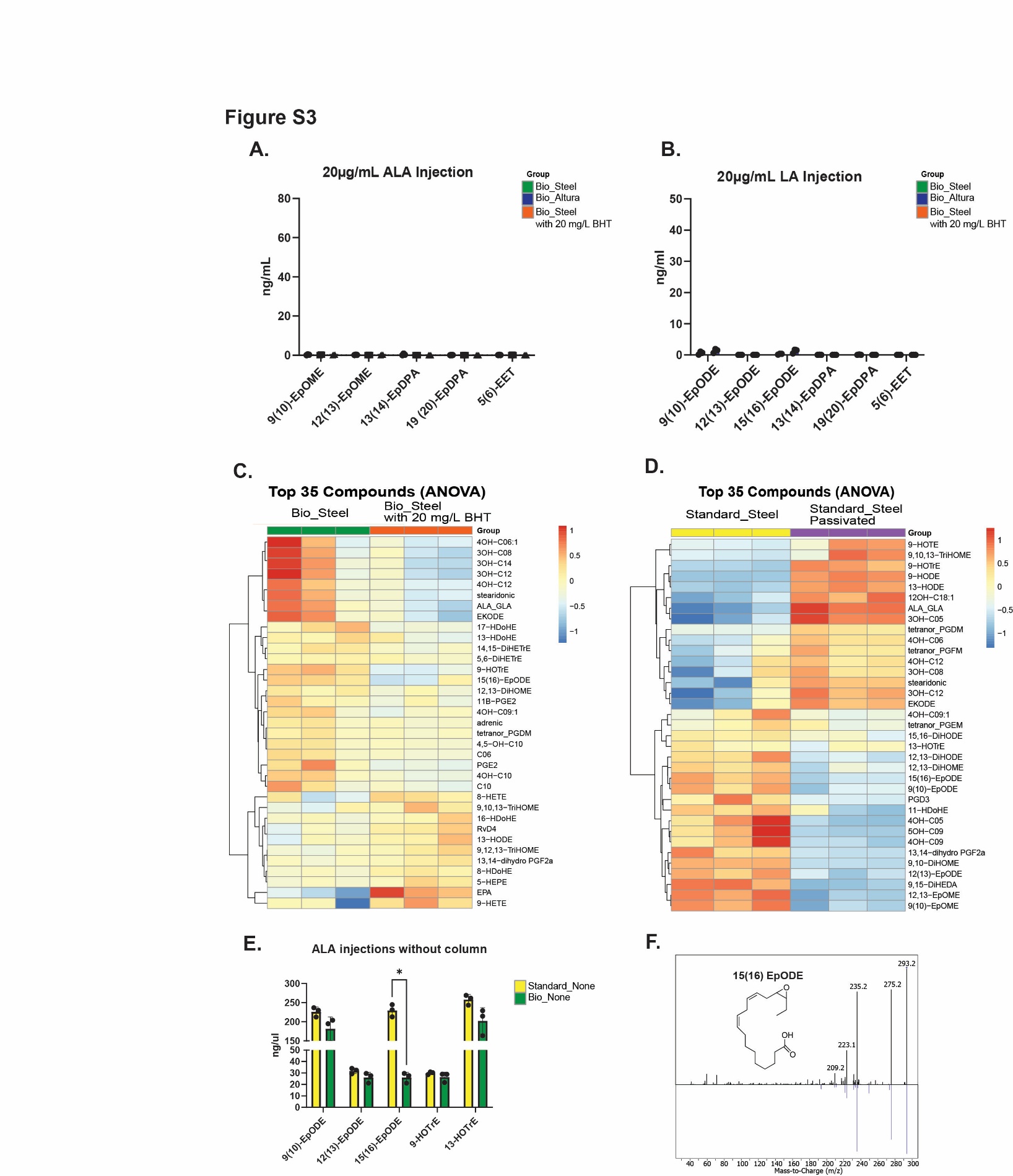

**Figure S3** PUFAs may be modified through oxidation in stainless-steel containing chromatography setups. (A) Epoxides from other PUFA precursors produced in 20 μg/mL ALA injection into various chromatography setups. (B) Epoxides from other PUFA precursors produced in 20 μg/mL LA injection into various chromatography setups. (C) Heatmap showing top 35 significantly changed oxylipins in healthy donor human plasma analyzed through the Bio_Steel setup with and without BHT added to mobile phase B. (D) Heatmap showing top 35 significantly changed oxylipins in human plasma in Standard_Steel before and after passivation. (E) Injections of ALA through unions (without chromatography columns) in standard vs. bio-inert LC instruments (F) Mass spectra of 15(16) EpODE generated inside Bio_Steel (top, black) vs. spectra of 15(16) EpODE previously generated (bottom, blue).

To further investigate the new method using Bio_Altura and blood fractionation, we studied standard separation in plasma fractionation, extraction recovery, and matrix effects in our setup. Standards were spiked into whole blood, and then blood was fractionated using the Ficoll-Paque extraction described in the main manuscript. We observed that the plasma fraction contained the majority of the standards spiked into whole blood, showing that blood fractionation is effective at sequestering extracellular oxylipins into the plasma fraction, but a small amount of standards may bind or become internalized in cells **(Figure S4a)**. Next, we tested the extraction efficiency of our method by comparing standards spiked into our system vs. an equal amount of standards that were taken through the extraction using plasma matrix for both groups. We observed a recovery level for most standards around 55%, which is lower than other lipidomics extractions, but consistent with previous reports using oxylipin solid-phase extraction**(Figure S4b)**. Finally, we tested the effects of matrix on standard relative intensity by spiking in standards to plasma matrix or methanol. We observed ion enhancement for most classes of standards, consistent with previous literature. **(Figure S4c)** We also evaluated CUDA as an extraction additive to determine whether soluble epoxide hydrolase inhibition during sample handling altered epoxide and diol oxylipin measurements. CUDA-containing extractions showed similar overall oxylipin profiles and technical reproducibility compared with the standard BHT-containing extraction condition, but did not broadly increase epoxide signals or decrease downstream diol signals **(Figure S4d)**. Therefore, CUDA was not incorporated into the final extraction method. To address method reproducibility, we added a technical precision analysis using plasma extracts. Across triplicate measurements, 70 analytes were detected, with a median CV of 4.1%; 67 of 70 detected analytes had CV values ≤20% **(Figure S4e).** We also evaluated short-term autosampler injection stability by reinjecting plasma extracts at 6 h and 12 h. Median responses were 98.9% and 94.2% of the initial plasma mean at 6 h and 12 h, respectively, indicating good short-term stability **(Figure S4f).** Inter-day precision was evaluated using pooled plasma QC samples analyzed in triplicate on three separate days, yielding a median between-day CV of 12.8%, with 73.4% of analytes meeting the ≤20% CV criterion **(Figure S4e).** These results demonstrate encouraging reproducibility across analytical days. This expands on **Figure 6** in the main manuscript.

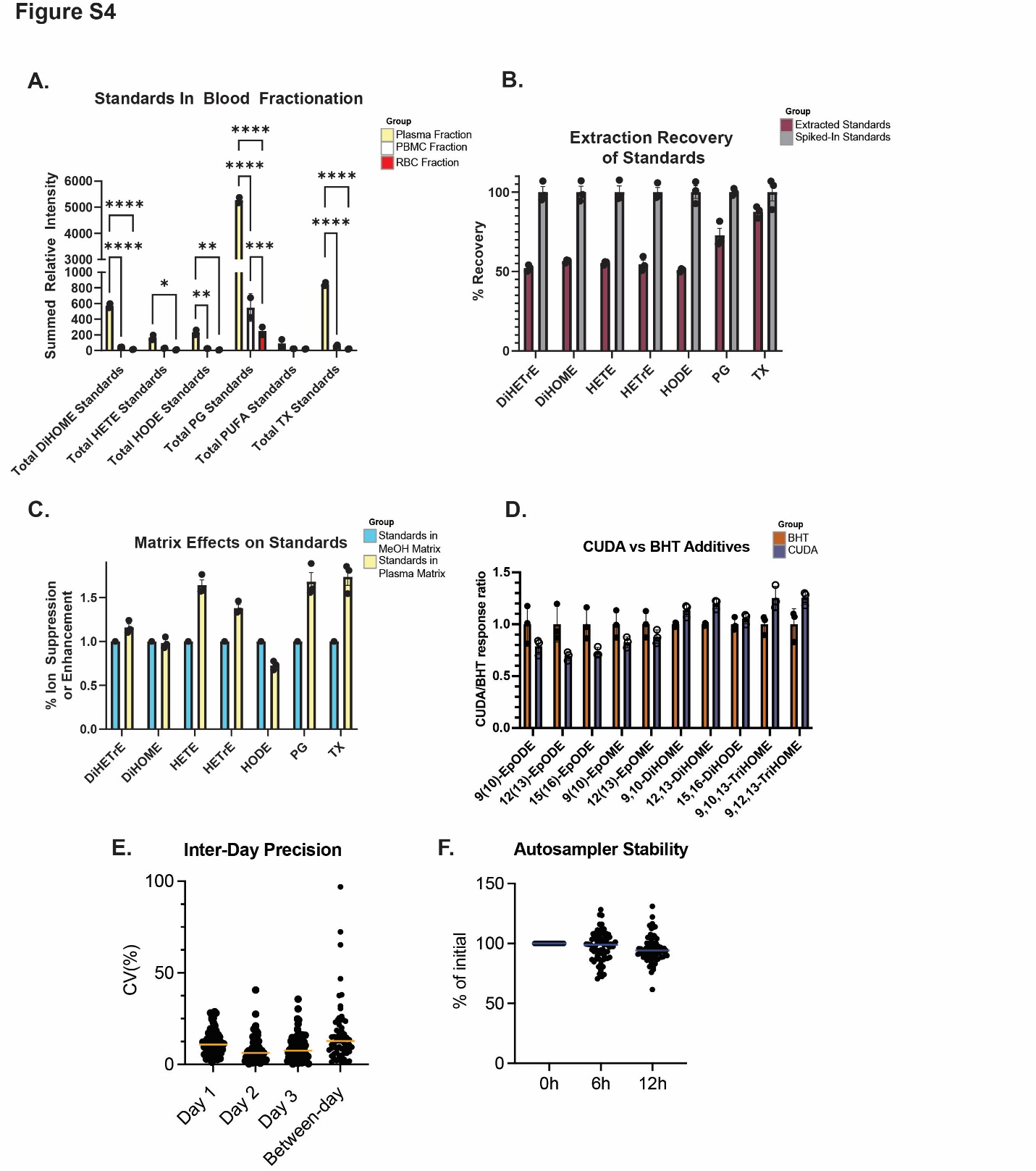

**Figure S4** Bio_Altura method investigation. (A) Bar graph showing levels of standards in each fraction of Ficoll-Paque fractionation after standards were spiked into whole blood before fractionation. (B) Bar graph showing extraction recovery of different standards, standards were summed by class. (C) Bar graph showing effects of plasma or methanol matrix on relative intensity of spiked-in standards. (D) Bar graph showing response ratios for BHT vs CUDA extraction additives. (E) Plot showing coefficient of variation of oxylipin species quantified for injections within the same day and across different days. (F) Plot showing % of oxylipins remaining after 6 and 12 hours inside the multisampler.

Using the Bio_Altura Setup, oxylipin analysis was performed on 6 additional human tissues to test the ability of the new instrumentation to detect oxylipins in a variety of tissues. **Table S1** lists the compounds detected in each tissue analyzed. Hydroxy fatty acids were also detected in tissues using this method. Islet analysis was limited by small tissue sample mass (10-20 mg), which was likely not optimal for analysis. Granulocytes, PBMCs, and RBCs were isolated from human plasma using Ficoll-Paque density gradient centrifugation. This expands on **Figure 6a** in the main manuscript.

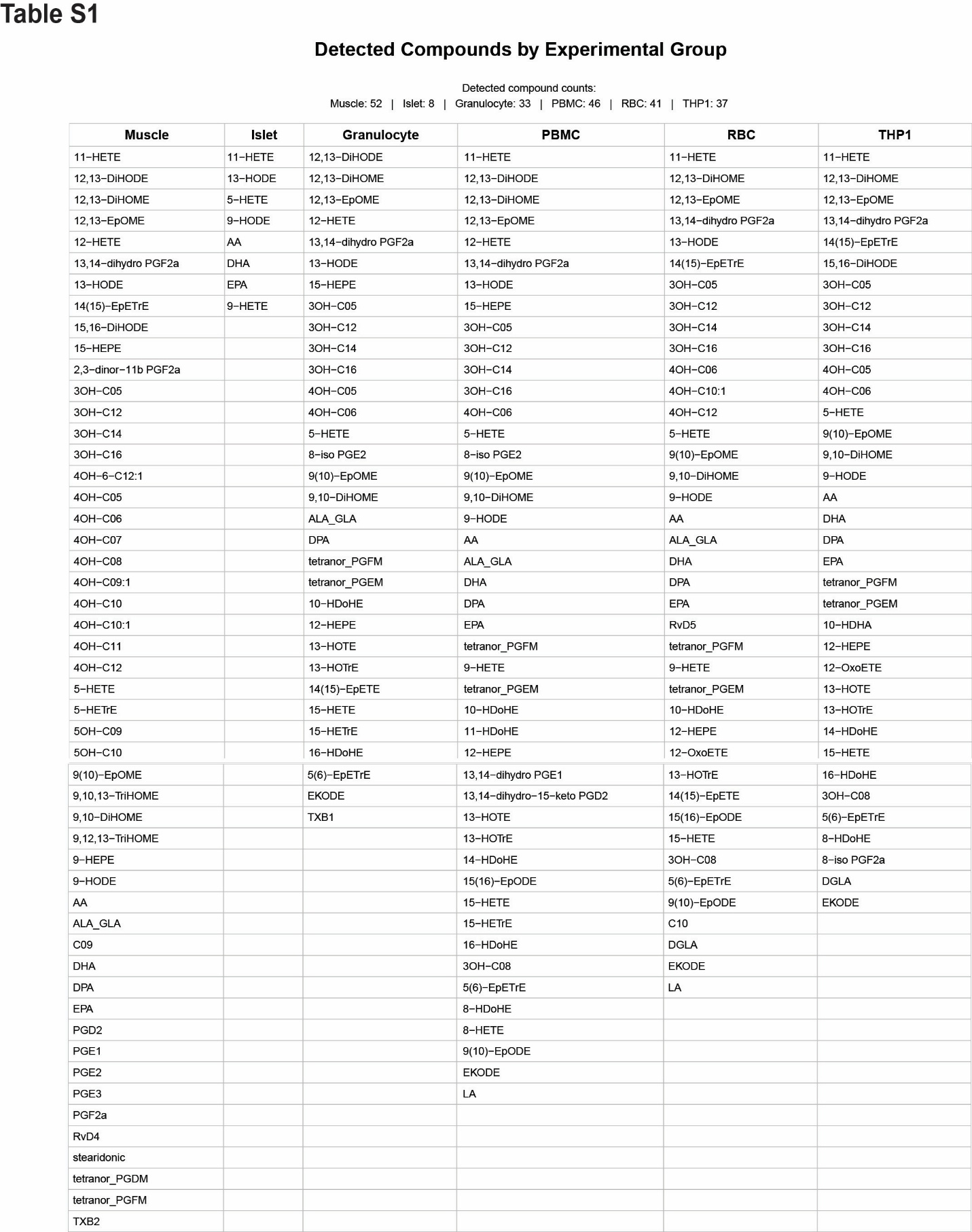

**Table S1**: Lists of compounds quantified using oxylipin analysis in 6 different human tissues. Tissues included skeletal muscle, islet cells, granulocytes, peripheral blood mononuclear cells (PBMCs), red blood cells (RBC), and THP-1 cells, a human monocyte-like cell line.

Specialized pro-resolving mediators were detected in two human tissues analyzed: RvD4 in muscle and Rvd5 in RBCs. While unexpected, these tissues could be the result of autoxidation, especially considering the RBC sample contains high amounts of iron and is exposed to ambient conditions for 30+ minutes due to the long duration of plasma fractionation using Ficolli Paque. The chromatograms provided in **Figure S5** do meet the 5:1 signal:noise ratio and achieve at least 10 points along the peak for quantification, as well as their standards. This expands upon the data seen in **Figure 6** of the main manuscript.

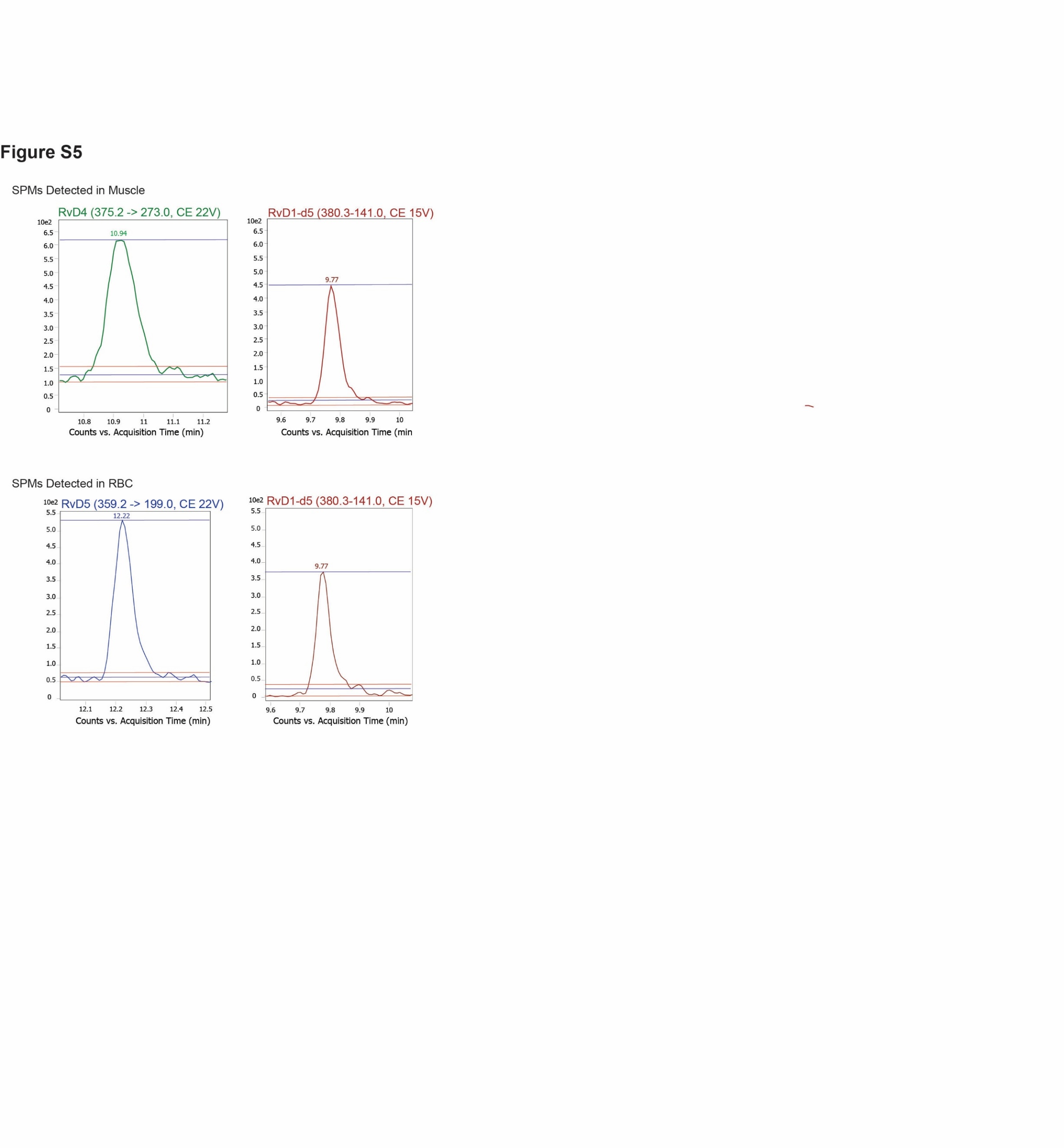
**Figure S5:** Chromatograms of all detected SPMs (2 total, one in muscle, one in RBC) alongside deuterated internal standard. MS/MS transitions, collision energy, and retention time are listed in each respective graph. Signal to noise is annotated using red lines for noise and blue for signal.

To determine separation ability of the chromatography system, we follow previously published guidelines and analyzed species that are markers of separation ability. **Figure S6** demonstrates good separation efficiency in various classes of DiHODE, HpETE, HDoHE, and prostaglandins.

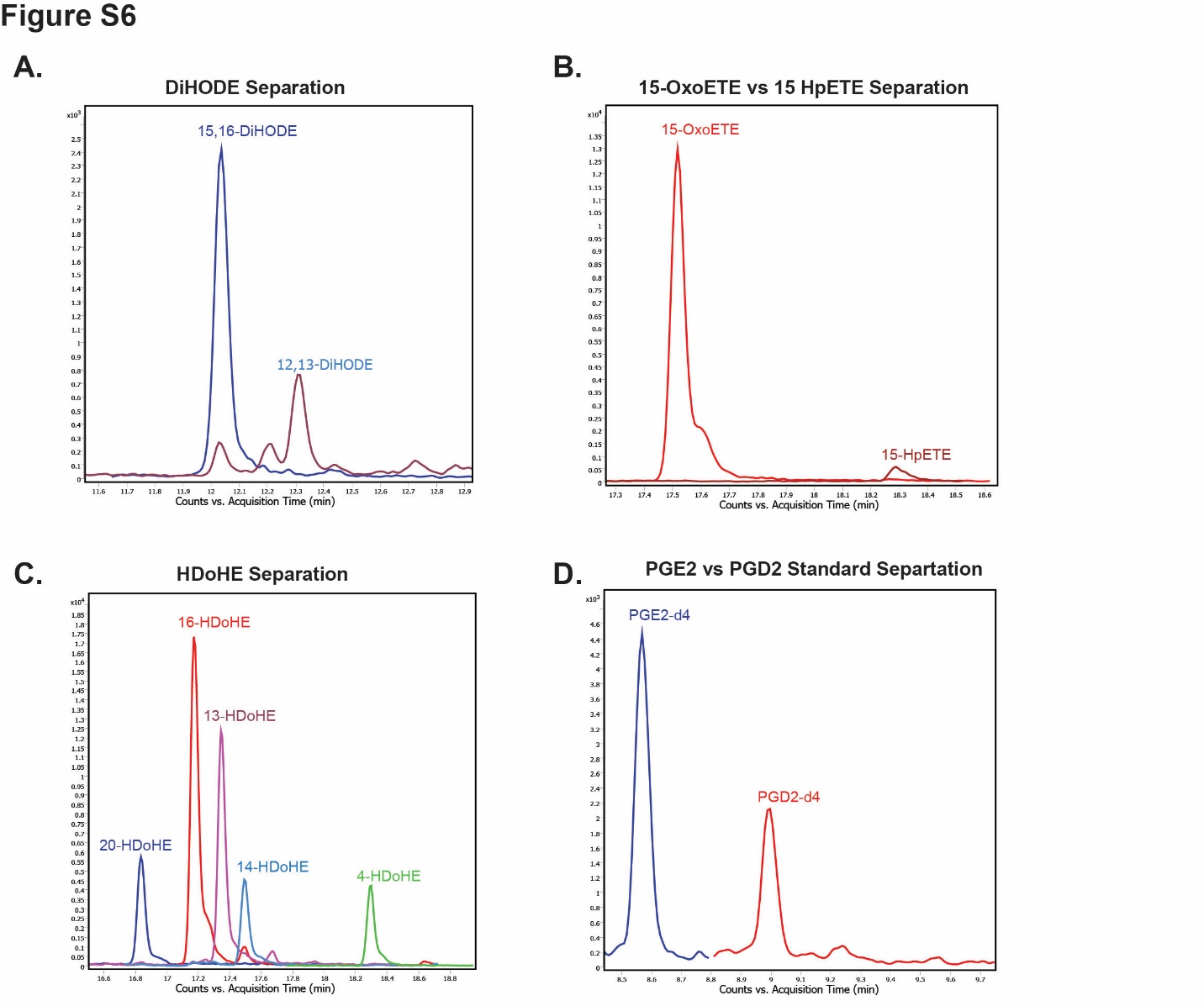
**Figure S6:** Separation ability of the chromatography system for oxylipin analysis. (A) Chromatograms for 15,16-DiHODE vs 12,13-DiHODE. (B) Chromatograms for 15-OxoETE vs 15-HpETE. (C) Chromatograms for various HDoHEs. (D) Chromatograms for PGE2-d4 vs PGD2-d4.

For analysis of compounds, relative quantification was used. Standards were paired to compounds based on similarity and used for analysis, the pairings are seen in **Table S2.** This expands on **Figure 6** of the main manuscript.

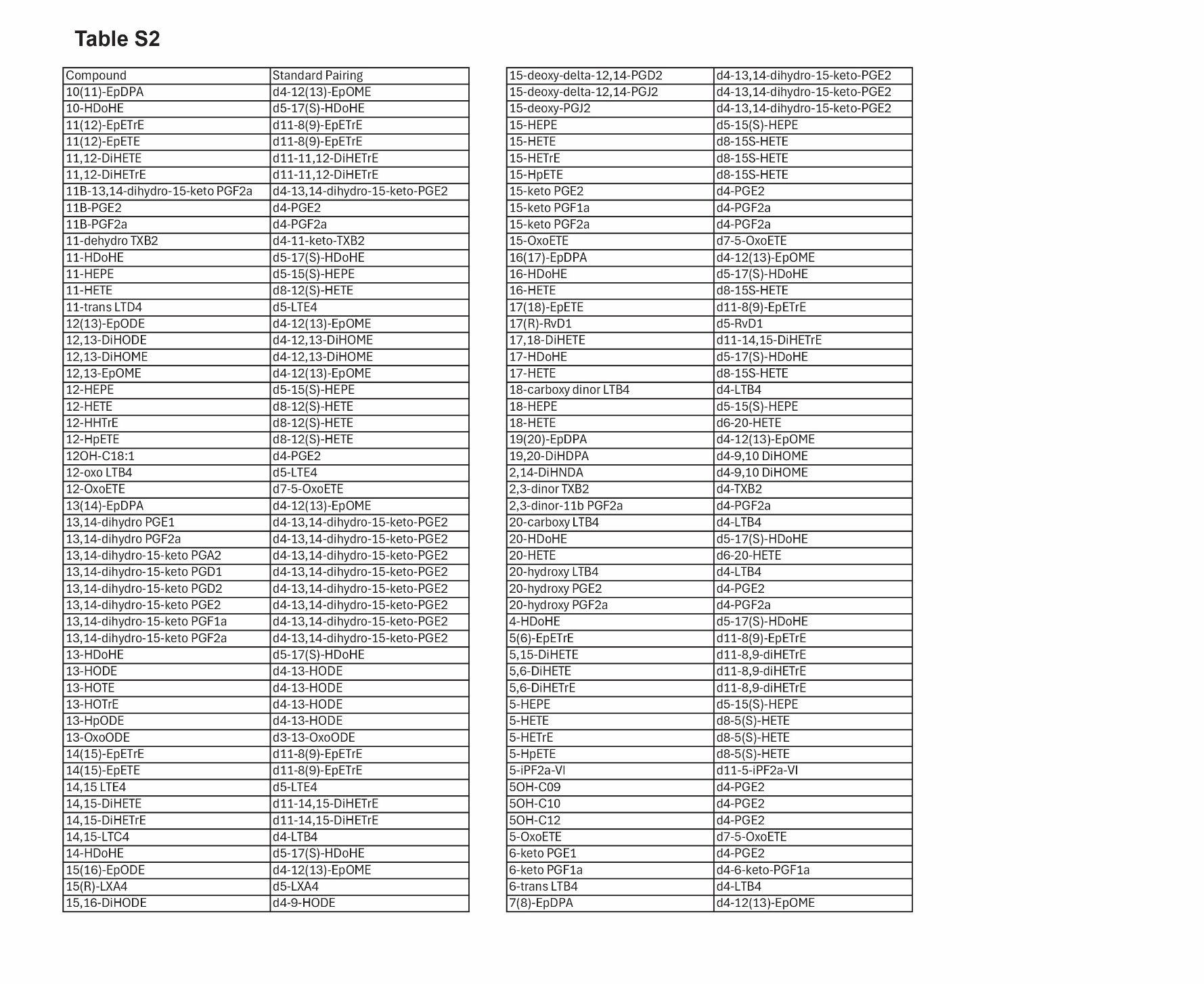

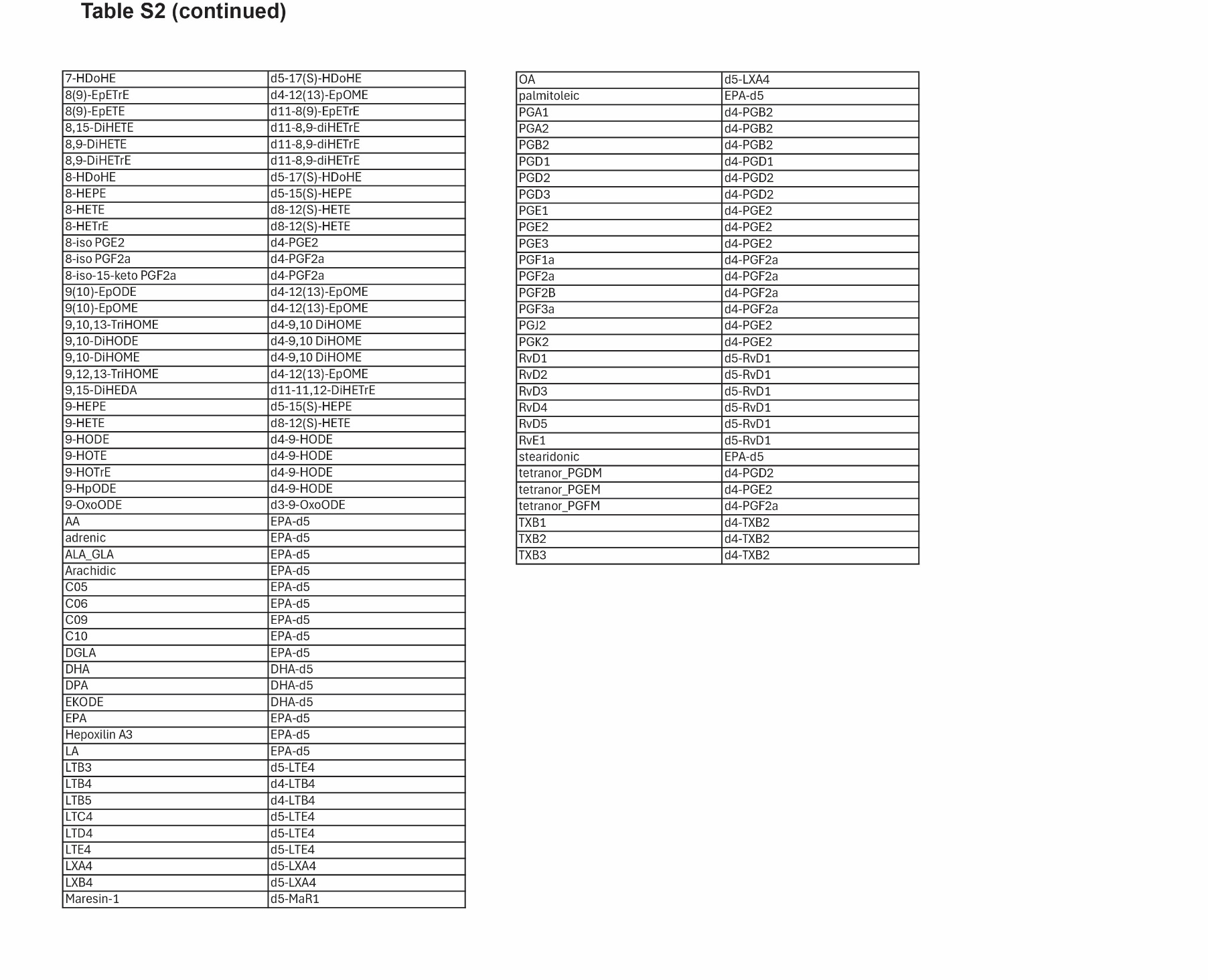

**Table S2:** Standard-compound pairings. Compounds of interest are listed on the left, standards used for quantification analysis for each compound are listed on the right
